# Compartmentalization of intestinal bacteria by hepatic ILC3s prevents infections after surgery

**DOI:** 10.1101/773150

**Authors:** Manuel O. Jakob, Daniel Sanchez-Taltavull, Bahtiyar Yilmaz, Thomas Malinka, Catherine Mooser, Spari Daniel, Lilian A. Salm, Katrin Freiburghaus, David Pereyra, Siegfried Hapfelmeier, Mojgan Masoodi, Patrick Starlinger, Deborah Stroka, Franziska Tschan, Daniel Candinas, Mercedes Gomez de Agüero, Guido Beldi

## Abstract

Infections after surgical interventions are assumed to be caused by contamination. We show by analyzing multicentric data of 6561 patients that surgical infections as well as sepsis had a predominantly enteric microbial signature irrespective of the type of surgery, suggesting failure of intestinal bacterial compartmentalization. In mice, we reveal that hepatic surgery induced dysregulation of intestinal and hepatic type 3 innate lymphoid cells (ILC3s) and intestinal leakage resulting in enteric bacterial translocation via lymphatic vessels. In the absence of hepatic ILC3s, inflammasome activation and the induction of antimicrobial peptide encoding genes, bacteria colonized remote systemic organs and impaired surgical outcomes. Conversely, mammalian-microbial commensalism is required for the education of host immunity to ensure optimal hepatic healing responses. In fact, microbial-derived products were sufficient for the induction of proliferative transcriptional networks in the mouse liver, as illustrated by serum transfer experiments, mass spectrometry and RNA expression analysis, indicating that the balanced exposure of the host to commensals is essential for recovery. This study reveals the intestinal origin of microbes causing complications after surgical interventions and highlights host protective mechanisms of controlled commensalism that prevent infections.

**One Sentence Summary:** Intestinal bacteria cause surgical infections

## INTRODUCTION

In Western countries, more than 20,000 surgical procedures are performed per 100,000 inhabitants annually, and infections are the most frequent post-surgical complication(*1*). The prevalence of healthcare-associated infections ranges between 3 and 6% among all patients, surgical site infections (SSI) being the most frequent with 29%(*2*). Infections occur locally as SSI or remotely, such as postoperative pulmonary infections. Both local and remote infections have the potential to end in sepsis(*3*). The World Health Organization (WHO) recently recognized sepsis as a global health priority and advocated for adherence to hygiene measures(*4*). These guidelines on hygiene are based on the assumption that microbes contaminating the operative field cause surgery-associated infections(*3-5*). However, the phylogenetic signature of bacteria isolated from surgical sites, draining lymph nodes, and from sites distant from the operative field includes largely enteric bacterial species that may suggest an intestinal origin for such infections(*2, 6, 7*).

In the intestine, barriers that limit the penetration of commensal and pathogenic microbes(*8*) consist of mucus, immunoglobulins, antimicrobial peptides, epithelial cells and immune cells(*9*). In particular, type 3 innate lymphoid cells (ILC3s), which are constitutively present at barrier sites and depend on the transcription factor Rorγt for their differentiation and function, play a pivotal role in host-microbial compartmentalization in the intestine and lung(*10, 11*). ILC3s have the specialized capability to sense both host-derived and environmental signals and can integrate and relay the cues to other cells through cytokine secretion or cell-cell interactions (*12*). Activated ILC3s secrete a polarized set of cytokines that include interleukin (IL)-17, IL-22, LTα1β2, and GM-CSF. In this way, ILC3s control phagocytosis, epithelial survival, and antimicrobial peptide secretion to combat infection, inflammation and to restore tissue homeostasis(*13*). The liver is a second and systemic barrier against bacterial infiltration that scans and clears circulating antigens and bacteria(*14, 15*). The steady-state hepatic innate immune system is composed of mainly Kupffer cells, natural killer (NK) cells, natural killer T (NKT) cells and ILCs. In the human liver, NKp44^-^ ILC3s are the dominant ILC population whose function remains largely unexplored(*16, 17*). Similar to that in the intestinal and pulmonary niches, ILC3-targeted cytokine signaling is relevant for hepatic tissue homeostasis(*18, 19*).

Here, we tested whether surgery disrupts the ILC3-mediated intestinal firewall and increases the susceptibility of mice to systemic infections triggered by the intestinal microbiota. By using large surgical patient cohorts, immune deficient and axenic mouse models, we sought to determine the source and the relevance of SSI. We found an intestinal origin of SSI that is the consequence of a dysregulation of intestinal and hepatic ILC3s. However, the microbiota has also been shown to be critical for the development of the host immune system(*20*). Such findings would support attempts in clinical practice to decrease the bacterial biomass in the intestine with the aim of preventing infections.

## RESULTS

### Microbes with an intestinal signature are a major cause of surgical site infection in patients

Infectious organisms after surgery are thought to originate from the surrounding skin or the operative field. To identify an association of the phylogenetic composition of SSI with the type of surgery, a prospective, multicenter cohort of patients with SSI as an endpoint was analyzed. SSI was observed to be the most frequent complication occurring in 8.6% (566 of 6561 patients) of all patients and up to 18.0% of patients undergoing high-risk (esophageal, hepatic, pancreatic, splenic and enteric) surgery. The microbial fingerprint of available SSI cultures (3515 patients) was analyzed using mass-spectrometry. Therefore, surgeries were grouped according to the site of surgery in intestinal surgeries opening the intestine (intestinal surgery), abdominal surgery without opening the intestine (non-intestinal surgery) and extra-abdominal surgery. The composition of bacteria did not vary between the different types of surgeries (Fig. 1a). Culturable bacteria found in SSI were mainly *Escherichia coli* and other Enterobacteriaceae (Fig. 1a), suggesting that infections may not primarily be the consequence of contamination and may be triggered via other routes of bacterial trafficking. In addition, the presence of infection but not the type of surgery dictated clinical consequences such as length of hospital stay (Fig. 1b) and resulted in increased mortality by 10-fold (Fig. 1c). Next, we aimed to determine the association of skin and intestinal microbial taxa with organisms found in SSI. Given the dominance of Firmicutes and Bacteroides phyla in the feces and Actinomyces and Fusobacterium dominating the skin, as assessed by 16S rDNA (Fig 1d), we applied routine clinical mass spectrometry of the bacteria cultured from skin and feces from 10 healthy volunteers with different ethnicities to determine the cutaneous and fecal microbial signatures at the genus level. In all specimens analyzed, *E. coli, Enterococcus spp*., and *Enterobacter spp*. were found only in the fecal samples, whereas *Staphylococcus spp*. and *Streptococcus spp.* were found only on the skin (Fig. 1e). Bacteria found in SSI (mainly *E. coli* and other Enterobacteriaceae) identified in the patient’cohort did mainly assign to bacteria that are more abundant in fecal samples suggesting that the majority of SSI have an intestinal origin (Fig. 1e).

**Fig. 1.**
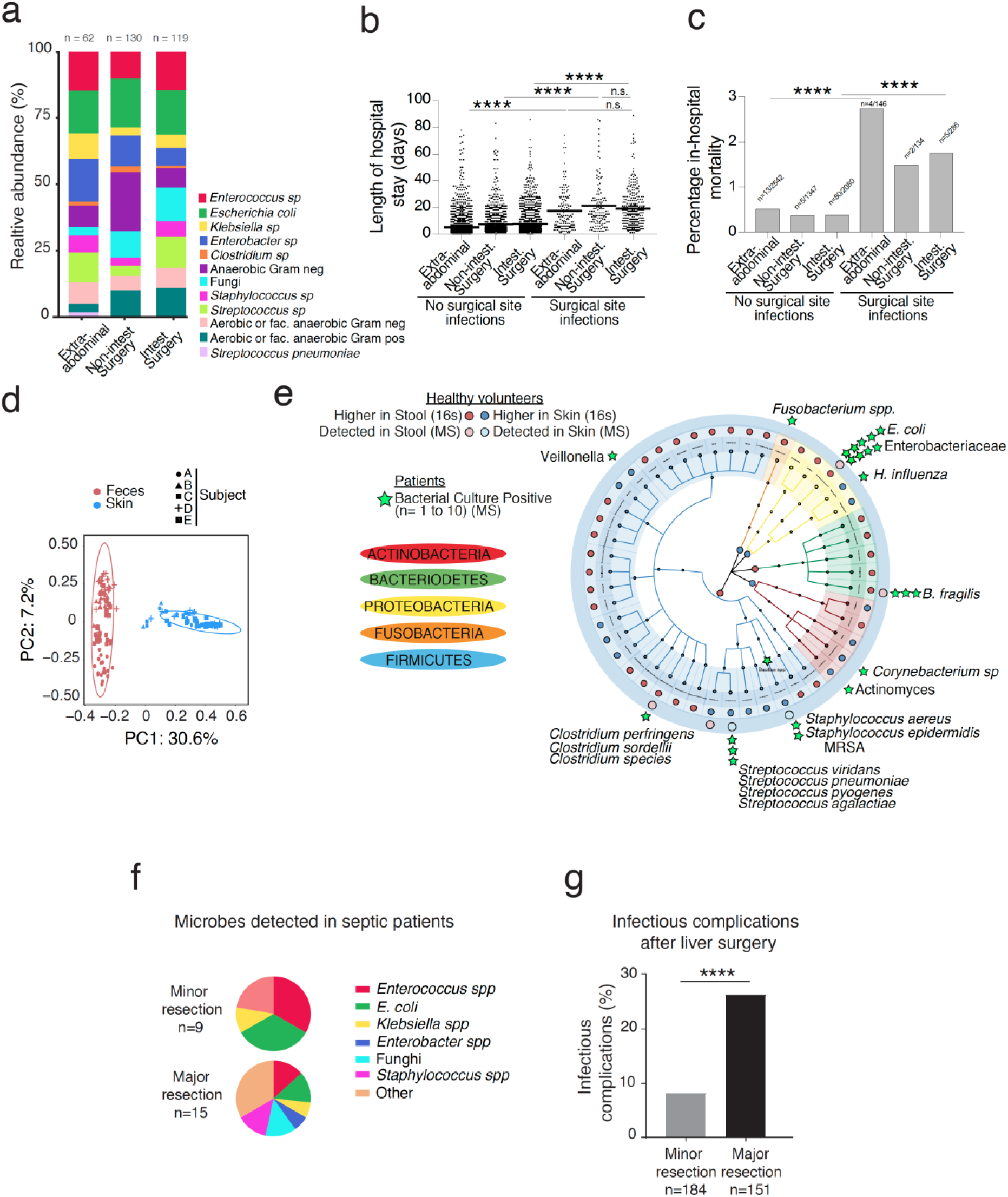
Bacteria cultured from surgical site infections have an intestinal signature. A multicenter cohort of 6561 patients from three Swiss surgical departments was analyzed. **(a, b, c)** Surgical site infections were grouped according to the site of main surgery in procedures that opened the intestine (intestinal surgeries - local spillage possible); abdominal surgery without opening the intestine (non-intestinal surgery) and extra-abdominal surgery (local spillage unlikely). **(a)** Relative abundance of bacterial species in SSI cultures obtained from 3515 patients that underwent extra-abdominal, non-intestinal and intestinal surgery. (**b**) Length of hospital stay on the y-axis and site of surgery on the x-axis for patients with or without surgical site infections. Each dot on the plot represents one patient. The line represents the median (n=4949 patients). (**c**) Mortality at 30 days post-surgery on the y-axis and site of surgery on the x-axis for patients with or without surgical site infections (n=6535 patients). **(d)** Microbial clustering is shown based on Bray-Curtis dissimilarity principal coordinate analysis (PCoA) metrics for longitudinally collected fecal and skin samples from 5 healthy individuals from 2 families. Non-parametric analysis of variance (Adonis) was used to test for significant differences between groups on the PCoA plot (*p*≤0.01). In this case, *p*<0.01 was considered significant. Ellipsoids represent a 95% confidence interval surrounding each disease group. **(e)** Overall representation of taxa significantly associated with either fecal or skin samples from the subjects analyzed in (d) are plotted and overlapped with bacterial strains that were identified with MALDI-TOF after culturing of the “fecal” or “skin” samples from ten different healthy volunteers. Statistical analysis for the taxonomy results was performed using the MaAsLin pipeline with BH-FDR (q value) and plotted using GraPhlAn. The red closed circle shows taxa with increased abundances in fecal samples, and the blue closed circle shows taxa with increased abundances in the skin samples. Stars on top of the graph represent the identified culturable bacteria in SSI in patients. One star correspond to 1-10 patients. **(f, g)** A second cohort of 335 patients undergoing minor or major liver surgery for colorectal liver metastasis was analyzed. **(f)** Identification of bacteria cultured from blood of patients with sepsis. **(g)** Frequency of infectious complications in patients that underwent minor or major liver surgery. The *p*-values are indicated as follows: \**p*≤0.05; ** *p*≤0.01; *** *p*≤0.001; **** p≤0.0001.

To investigate the potential contribution of the extent of surgery in contrast to the procedure itself as a cause of enteric microbial translocation to systemic sites, a second cohort of 335 patients undergoing major or minor liver surgery was analyzed. As in the previous cohort, analysis of patients who suffered from sepsis after minor and major hepatic surgery revealed that the circulating bacteria had an intestinal microbial signature (Fig. 1f). The frequency of infectious complications (SSI, sepsis) was higher in patients who underwent major liver resections (Fig. 1g) excluding the procedure as a cause for contamination and suggesting that the extent of the surgical trauma may disturb the intestinal integrity increasing bacterial translocation. These clinical data further support the hypothesis that microbes causing SSI and sepsis are largely of intestinal origin and that the extent of surgery, rather than the site of surgery, dictates the occurrence of an SSI.

### Major liver surgery provokes intestinal microbial translocation under ILC deficiency

To understand the impact of surgery on systemic dissemination of intestinal bacteria and the potential contribution of immune cell populations, we modeled major human surgery using a two-third partial hepatectomy in C57BL/6 wild-type mice (Fig. 2a). Twenty-four hours after liver surgery, bacteria were detected in mesenteric lymph nodes (MLNs), whereas the liver remained sterile and exhibited optimal regeneration at 48 hours post-surgery (Fig. 2b, c)(*21, 22*). Since compartmentalization of intestinal microbiota in the gut lumen under homeostatic conditions depends on functional intestinal ILC3s(*10, 23*), we investigated the impact of liver surgery on ILC3s and other immune cell populations. Cellular fractions of MLNs, the *lamina propria* in the small intestine and colon and from the liver were analyzed by flow cytometry following partial hepatectomy (Fig. S1). The frequency of small intestine and hepatic ILC3s but not from the MLN and colon was affected by partial hepatectomy, whereas NK cell, T cell, ILC1 and ILC2 populations remained unchanged in the different tissues analyzed (Fig. 2d,e and Fig. S2). At 24 hours post-surgery, the small intestinal *lamina propria* CD3^-^ CD127^+^Rorγt^+^ ILC3 population significantly decreased (Fig. 2d) in parallel with the level of the effector cytokine IL-17, but not IL-22 (Fig. S3a, b). In contrast to the decrease observed in the small intestine, the hepatic CD3^-^CD127^+^Rorγt^+^ ILC3 population expanded after partial hepatectomy at 24 hours post-surgery (Fig. 2e). The decrease in small intestinal ILC3 and expansion in hepatic ILC3 population (Fig. 2d,e) was not observed in germ-free wild-type mice that underwent partial hepatectomy, indicating the requirement of microbiota for changes in the ILC3 population (Fig. S3c,d)(*16*).

**Fig. 2.**
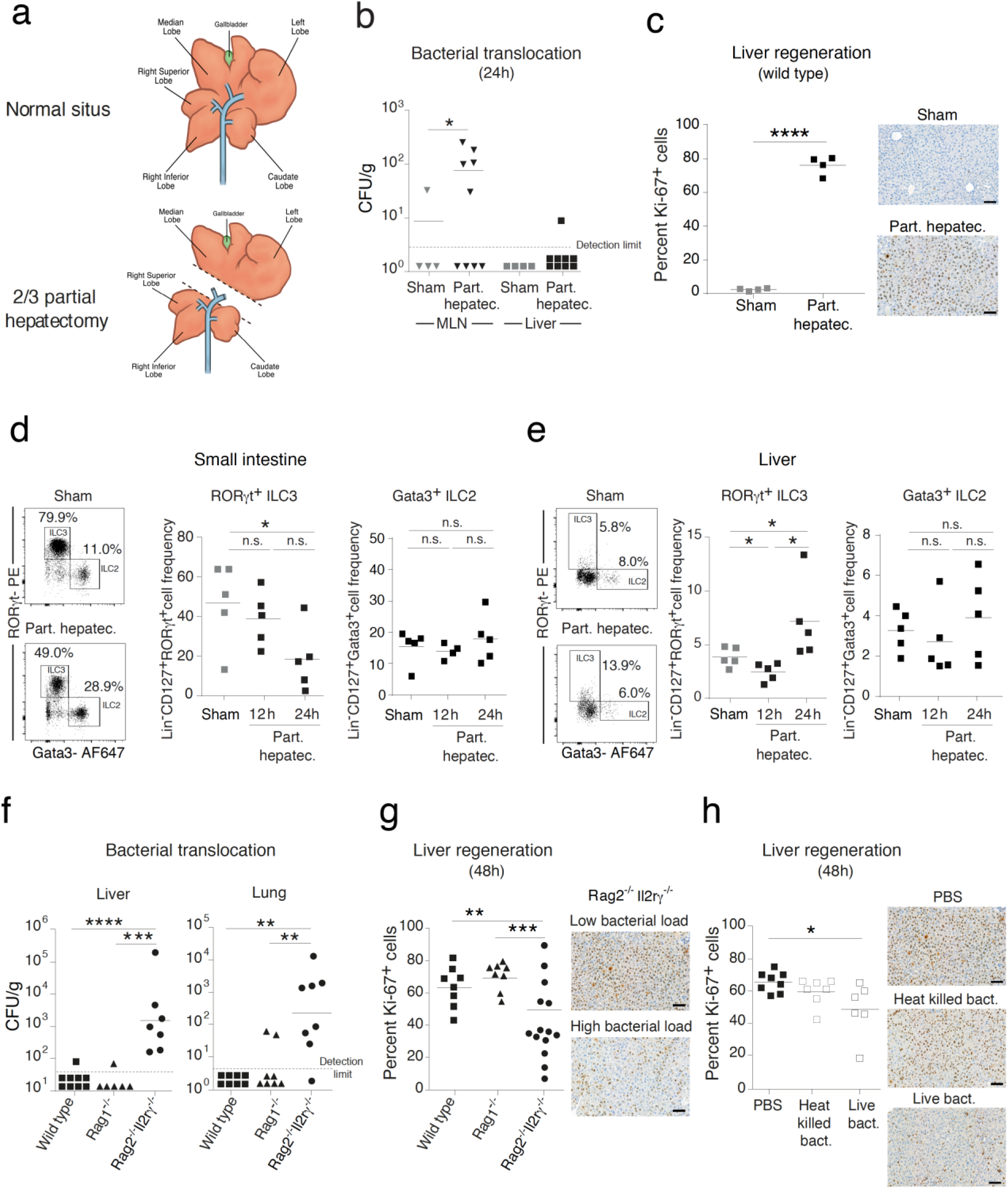
Type 3 innate lymphoid cells (ILC3s) control the systemic penetration of intestinal bacteria. Specific pathogen-free (SPF) wild-type mice **(a-e, h)** or SPF wild-type, *Rag1*^*-/-*^ and *Rag2*^*-/-*^*Il2rγ*^*-/-*^ mice **(f, g)** underwent major surgery modeled by a two-thirds partial hepatectomy. **(a)** Schematic representation of two-thirds partial hepatectomy in mice in which the left and median liver lobe are removed. **(b)** Bacterial titers in mesenteric lymph nodes (MLNs) and liver 24 hours after partial hepatectomy or sham surgery. **(c)** Liver regeneration was assessed by estimating the percentage of Ki-67-positive hepatocytes 48 hours after partial hepatectomy or sham surgery. Representative images are presented on the right and quantification of proliferation is presented on the left. **(d, e)** Frequency of small intestinal *lamina propria* and liver Rorγt^+^ ILC3s gated as live^+^Lin^-^Gata3^int/-^Rorγt^+^ cells and Gata3^+^ ILC2s gated as live^+^Lin^-^Rorγt^-^Gata3^+^ 24 hours after partial hepatectomy or sham surgery. **(f)** Bacterial titers in the indicated organs and **(g)** the percentage of Ki-67-positive hepatocytes 48 hours after partial hepatectomy or sham surgery are shown. **(h)** The percentage of Ki-67-positive hepatocytes after intravenous injection of PBS, heat-killed or live *E. coli* K-12 strain JM83 into SPF wild-type mice before and after partial hepatectomy. The mice were analyzed 48 hours after partial hepatectomy. Representative images are presented on the right and quantification of proliferation is presented on the left (h, i). Scale bars (g, h): 100 μm. Normalized values were analyzed by Student’s t test to compare two experimental groups or by ANOVA to compare more than two groups in parallel. Data are representative of 4 (b, c), or 2-3 (d-h) independent experiments (8-15 mice per group). (g) Data pooled from 3 independent experiments. The *p*-values are indicated as follows: \**p*≤0.05, ** *p*≤0.01, \*\*\**p*≤0.001, \*\*\*\**p*≤0.0001.

To study the contribution of ILC3s to the post-surgical infection in the context of liver surgery, we compared the source of infection following partial hepatectomy in *Rag2*^*-/-*^*Il2rγ*^*-/-*^ mice which lack ILC, natural killer, NKT, T and B cells to *Rag1*^-/-^ mice which lack only B, T and NKT cells and to C57BL/6 wild-type mice. Bacteria translocate to MLN in wild-type, *Rag1*^-/-^ and *Rag2*^*-/-*^*Il2rγ*^*-/-*^ mice (Fig S4a). In *Rag2*^*-/-*^*Il2rγ*^*-/-*^ mice, bacteria were detected in the liver, lung (Fig. 2f) and spleen, whereas these organs in C57BL/6 wild-type and *Rag1*^-/-^ mice remained sterile which points to the importance of the innate lymphoid compartment, including ILC and natural killer cells in barrier protection (Fig. 2f and Fig. S4b-e). This was not dependent on the intestinal microbial composition given that the bacterial species identified in the liver, spleens, MLNs and lungs of *Rag2*^*-/-*^*Il2rγ*^*-/-*^ mice mainly included *Escherichia coli, Enterococcus faecalis* and *Enterobacter hormaechei* and to a lesser abundance *Staphylococci ss*, which are all members of the intestinal microbiota (Fig. S4f-i)(*24*). We next assessed the effect of intestinal bacterial dissemination on proliferative hepatocyte indices, which are the most relevant outcome parameters after hepatic surgery. Translocation of intestinal bacteria to the liver following partial hepatectomy in *Rag2*^*-/-*^*Il2rγ*^*-/-*^ mice (see Fig. 2f) was associated with failure of liver regeneration with mice with a higher bacterial load being associated with lower hepatocellular proliferation (Fig. 2g and Fig. S5a, b). To confirm the deleterious effect of circulating bacteria on hepatic regenerative responses, we intravenously injected *E. coli* JM83 into specific pathogen-free (SPF) wild-type mice before and after partial hepatectomy (Fig. S5c). In comparison to wild-type mice that received PBS or heat-killed bacteria, hepatocellular proliferation and expression of cell cycle regulatory genes (*Foxm1b and Ccna2*) in the liver were impaired in wild-type mice that received live bacteria (Fig. 2h and Fig. S5d) (*16*).

Considering the observed systemic microbial translocation in *Rag2*^*-/-*^*Il2rγ*^*-/-*^ mice (see Fig. 2f) that may be due to differences in the intestinal colonization status, we examined whether monocolonization with *E. coli* K-12 strain JM83 in germ-free mice recapitulate the effect observed in SPF mice. Therefore, we subjected germ-free wild-type, *Rag1*^-/-^ and *Rag2*^*-/-*^*Il2rγ*^*-/-*^ mice to bacterial colonization by intragastric inoculation of *E. coli* 12 hours prior to partial hepatectomy (Fig. S6a). We observed that a minor proportion of bacteria can penetrate the intestinal barrier and reach the liver in sham germ-free *Rag2*^*-/-*^*Il2rγ*^*-/-*^ mice, but not in sham germ-free wild-type or *Rag1*^-/-^ mice (Fig. S6b). Further, partial hepatectomy induced an intestinal leak in monocolonized *Rag2*^*-/-*^*Il2rγ*^*-/-*^ mice that led to a high incidence of sepsis-induced mortality (Fig. S6c), thereby pointing towards the role of ILC3s in host-microbial mutualism.

These experiments suggest that surgery directly disrupts compartmentalization of intestinal microbes illustrated by a decreased frequency and function of intestinal ILC3s. Failure to effectively contain intestinal bacteria results in systemic spread of bacteria infecting organs and eventually impairing outcome after surgery.

### Major surgery induces intestinal leakage that allows bacteria to penetrate from the small intestine, via the lymphatic vessels into the systemic circulation

In order to understand if metabolic or cytokine dysregulation induced by partial hepatectomy contributes to intestinal barrier disruption, spectrometry and cytokine array were performed. We detected marked alterations in metabolites and cytokines in the serum of wild-type mice eight hours after partial hepatectomy (Fig. 3a,b and Table S1, S2). The levels of metabolites associated with bile acid biosynthesis and the inflammatory cytokine IL-6 were significantly increased, both of which have been described to increase intestinal permeability and impair intestinal barrier function (Fig. 3c and Fig. S7) (*25, 26*). Fecal albumin, a sign of intestinal barrier leakage, was detected in wild-type, *Rag1*^-/-^ and *Rag2*^*-/-*^*Il2rγ*^*-/-*^ mice after surgery, although the concentration was highest in the feces of *Rag2*^*-/-*^*Il2rγ*^*-/-*^ mice (Fig. 3d), suggesting that epithelial leakage promoted by hepatic surgery is enhanced in the absence of ILC. To evaluate the penetration of bloodstream contents into the intestinal tissues, we investigated the state of the intestinal vascular system using *in vivo* confocal endomicroscopy (Cellvizio) in mice that were intravenously injected with a fluorescent dye (70 kDa FITC-dextran) after partial hepatectomy. We found increased fluorescence by partial hepatectomy in both, wild-type and *Rag2*^*-/-*^*Il2rγ*^*-/-*^ mice, indicating that surgery promotes endothelial leakage and the release of bloodstream contents into the *lamina propria* of the intestine (Fig. 3e and Movie S1,S2)(*27*), which may contribute to intestinal barrier disruption and bacterial penetration.

**Fig. 3.**
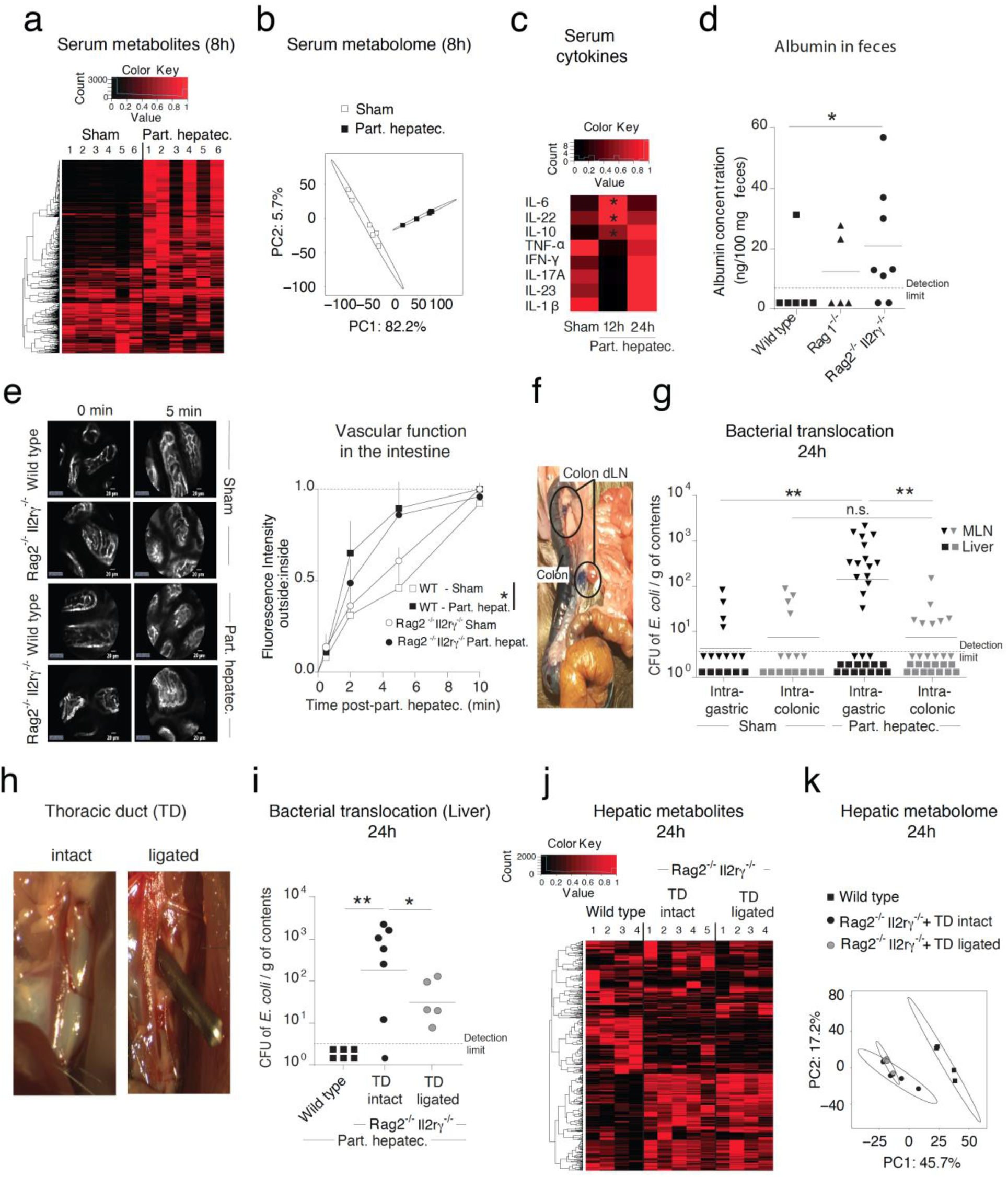
Translocating bacteria originate from the small intestine and traffic into the systemic circulation via the lymphatic vessels. Wild-type and *Rag2*^*-/-*^*Il2rγ*^*-/-*^ **(d, e, h-k)** mice underwent two-thirds partial hepatectomy. **(a)** Heatmap and **(b)** principal coordinates analysis (PCoA) of all metabolites identified in the serum 8 hours after partial hepatectomy analyzed by high-resolution mass spectrometry in positive ionization mode. **(c)** Heatmap of cytokines measured in the serum of SPF wild-type mice at the indicated time points (n ≥5 mice in each group, *p≤0.05 between sham and 12 hours partial hepatectomy). **(d)** Germ-free wild type, Rag1^-/-^ and *Rag2*^*-/-*^*Il2rγ*^*-/-*^ mice were short-term colonized by gavage with 10^10^ CFU of *E. coli* JM83 12 hours prior to partial hepatectomy, followed by measurement of fecal albumin by ELISA 24 hours post-surgery. **(e)** Immediately after partial hepatectomy, intestinal permeability was visualized in wild-type and *Rag2*^*-/-*^*Il2rγ*^*-/-*^ mice using endomicroscopy for 30 min after intravenous injection of 70 kDa FITC-dextran (Also see Movies S1 and S2). **(f)** Subserosal injection of Evans blue into the colon was performed to stain and visualize the draining lymph nodes. **(g)** SPF wild-type mice were challenged intragastrically or intracolonically with 10^10^ CFU of *E. coli* JM83 12 hours prior to partial hepatectomy or sham surgery. MLNs and livers collected 24 hours after the partial hepatectomy were minced and plated to estimate CFUs of translocation of *E. coli* JM83. **(h)** Imaging of thoracic duct ligation. The thoracic duct was clipped with a small surgical clip occluding lymphatic flow without disruption of the duct. **(i)** The thoracic duct in *Rag2*^*-/-*^*Il2rγ*^*-/-*^ mice was ligated, and these mice along with wild-type mice and *Rag2*^*-/-*^*Il2rγ*^*-/-*^ mice with intact thoracic duct were colonized intragastric with 10^10^ CFU of *E. coli* JM83. Twelve hours after colonization, the mice underwent partial hepatectomy or sham surgery, and liver samples were collected 24 hours after the operation, followed by enumeration of bacterial titers. **(j, k)** Following ligation of the thoracic duct, oral bacterial colonization for 24 hours and partial hepatectomy, mass spectrometry analysis was performed on whole liver tissue from *Rag2*^*-/-*^*Il2rγ*^*-/-*^ mice. Liver tissues from wild-type mice and *Rag2*^*-/-*^*Il2rγ*^*-/-*^ mice with intact thoracic duct that were subjected to bacterial colonization for 24 hours and partial hepatectomy were used as controls. Heatmap and PCoA of metabolites were identified in the positive ionization mode. The data shown are the arithmetic mean for the linear scale ±SD and the geometric mean for the logarithmic scale. Normalized values were analyzed by Student’s t test to compare two experimental groups or by ANOVA to compare more than two groups in parallel. Data are representative of 2-3 independent experiments or pooled from 2 independent experiments. Data for (d) are representative of three experiments. The *p*-values are indicated as follows: \**p*≤0.05; ** *p*≤0.01.

Since bacteria translocated from the intestinal lumen to MLNs (Fig. S4a), we sought to dissect the anatomical route of translocation. The lymphatic drainage from the intestinal lumen to MLNs is anatomically distinct in the colon compared to the small intestine (Fig. 3f)(*28, 29*). To identify the intestinal compartment from which bacteria translocate to the MLNs, we delivered *E. coli* JM83 to C57BL/6 wild-type mice using two routes – oral (intragastric) or rectal (intracolonic) prior to partial hepatectomy, and measured the enteric bacterial load in the local MLN and the liver. Significant bacterial load in the MLNs was observed with intragastric delivery, but not with intracolonic delivery, revealing that the primary site of translocation is the small intestine rather than the colon (Fig. 3g). Intragastric delivery of *E. coli* JM83 into C57BL/6 wild-type mice neither promoted the penetration of intestinal microbes to the liver nor altered liver regeneration (Fig. 3g and Fig. S8). The predominant translocation from the small intestine seems to be site-specific given that we observed slight tight junction gene expression changes in the colon in addition to a more marked mucosal proinflammatory cytokine immune response (Fig. S9). Potential routes for microbial trafficking from the MLN to the systemic circulation are lymphatic vessels and splanchnic blood circulation(*27, 30*). To determine the route of circulating microbes, the thoracic duct was ligated (Fig. 3h), followed by oral inoculation with *E. coli* JM83 and partial hepatectomy. *E. coli* JM83 failed to reach the liver in *Rag2*^*-/-*^*Il2rγ*^*-/-*^ mice with thoracic duct ligation (Fig. 3i). Thus, our results suggest that enteric bacteria preferentially transit from the small intestinal *lamina propria* using the lymphatic vessels and reach the liver by bypassing the splanchnic circulation.

Since systemic dissemination of intestinal bacteria was seen to impair surgical outcomes (see Fig. 2f,g and Fig. S5a,d), we hypothesized that hepatic metabolism would also be altered due to systemic bacterial dissemination. To test this hypothesis, non-targeted metabolic profiling was conducted on hepatic tissue from wild-type mice and from *Rag2*^*-/-*^*Il2rγ*^*-/-*^ mice with or without ligation of the thoracic duct and orally inoculated with *E. coli* JM83 prior to partial hepatectomy. Metabolic profiling revealed marked changes in specific metabolic pathways in *Rag2*^*-/-*^*Il2rγ*^*-/-*^ compared to wild-type control mice after thoracic duct ligation (Fig. 3j and Fig. S10a,b and Table S3). In particular, wild-type mice that display complete hepatic regeneration unlike *Rag2*^*-/-*^*Il2rγ*^*-/-*^ mice, exhibited elevated carnitine shuttle metabolism, which is consistent with the association of fatty acid metabolism with hepatic regeneration (Fig. S10c)(*31-33*). In *Rag2*^*-/-*^*Il2rγ*^*-/-*^ mice, no difference in the metabolome was found after blockade of the lymphatic drainage (Fig. 3j,k and Fig. S10a,b), suggesting that metabolic alterations in the liver are independent of intestinal bacteria trafficking through the thoracic duct.

### Inflammasome-controlled hepatic ILC3s maintain systemic antibacterial defense

The systemic dissemination of intestinal bacteria in *Rag2*^*-/-*^*Il2rγ*^*-/-*^ mice (see Fig. 2f) highlights the role of the innate lymphoid compartment in host protection. Since ILC3s compartmentalize bacteria via the production of antimicrobial peptides(*10*), we examined the expression of antimicrobial peptide-encoding genes in the livers of SPF (Fig. 3a) and germ-free (Fig. S11a) wild-type mice that underwent a two-thirds partial hepatectomy. We observed that the presence of microbes resulted in an increase of the expression of antimicrobial peptide-encoding genes specifically in the livers of wild-type but not germ-free mice at 24 hours (Fig. 4a and Fig. S11a). Such changes were not observed in the intestine (Fig. S11b, c). These results suggest that antimicrobial peptides are used to clear systemic bacteria(*34*).

**Fig. 4.**
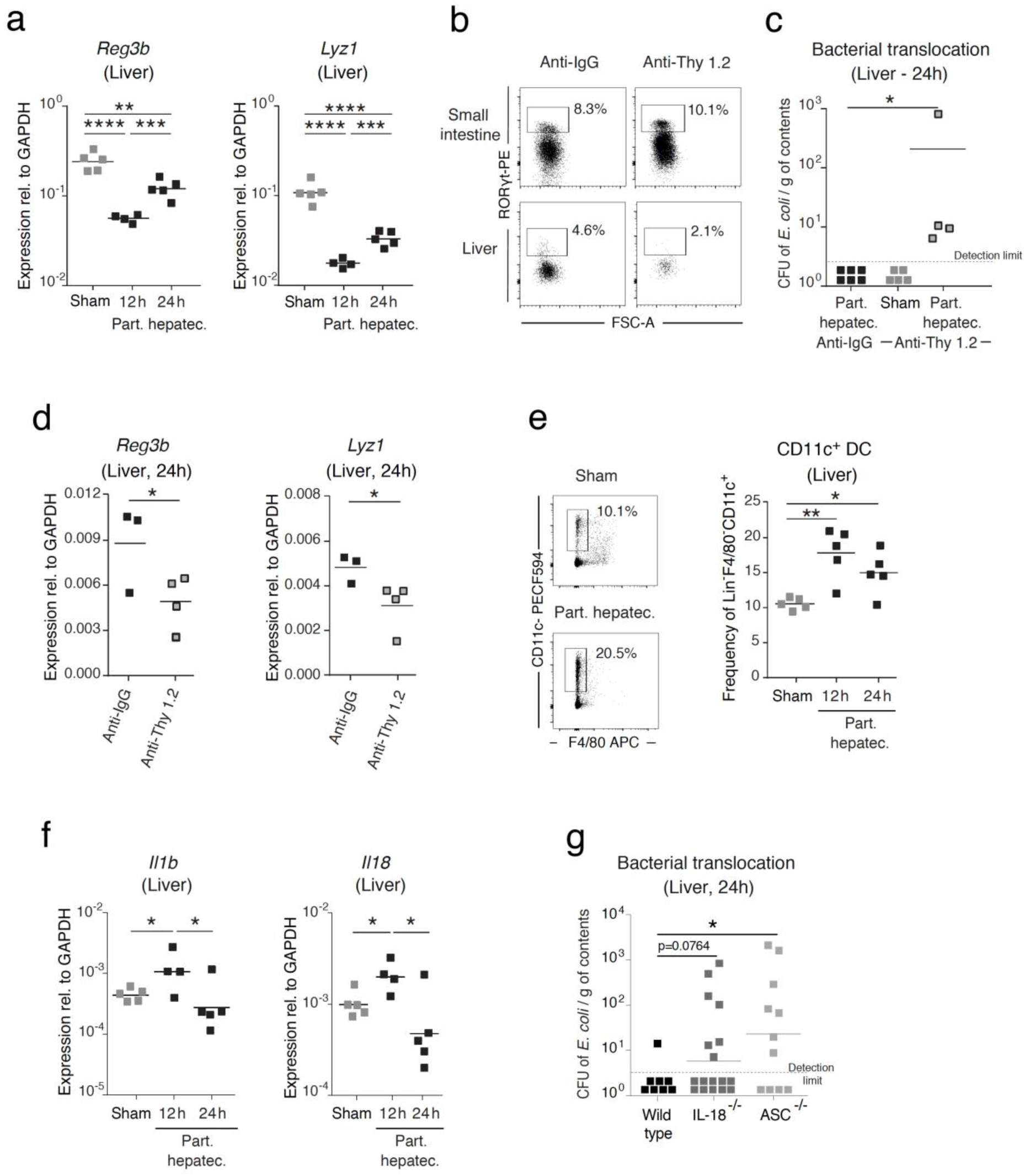
Systemic infections are controlled by the inflammasome, hepatic ILC3s and via antimicrobial peptides. SPF wild-type mice **(a-f)** or SPF wild-type, IL-18^-/-^ and ASC^-/-^ mice (**g**) were subjected to a two-thirds partial hepatectomy. **(a)** Whole tissue RNA was isolated from the liver of mice that were sham-operated or subjected to partial hepatectomy, and the expression of antimicrobial peptide-encoding genes was analyzed by RT-qPCR at the indicated time points. Gene expression was normalized to *GAPDH*. **(b)** Frequency of hepatic Rorγt^+^ ILC3s gated as live^+^Lin^-^Gata3^-^Rorγt^+^ cells after treatment with an anti-Thy 1.2 antibody or anti-IgG injected intraperitoneally. **(c)** Bacterial titers of *E. coli* JM83 in the liver following treatment with anti-Thy1.2 antibody or anti-IgG and challenge with 10^10^ CFU of *E. coli* JM83 12 hours prior to partial hepatectomy or sham surgery. **(d)** Expression of antimicrobial peptide-encoding genes was analyzed by RT-qPCR in whole liver tissue following treatment with anti-Thy1.2 antibody or anti-IgG and challenge with 10^10^ CFU of *E. coli* JM83 12 hours prior to partial hepatectomy. **(e)** Flow cytometry analysis of hepatic dendritic cells gated as live^+^CD3^-^ CD19^-^Ly6C^-^Ly6G^-^SiglecF^-^F4/80^-^CD11c^+^ cells was performed in sham-operated mice or at the indicated time points after partial hepatectomy. **(f)** Expression of *IL-1β* and *IL-18* was analyzed in whole liver tissue in sham-operated mice or at the indicated time points after partial hepatectomy using RT-qPCR. Gene expression was normalized to *GAPDH*. **(g)** Bacterial titers in the liver were quantified in SPF wild-type, IL-18^-/-^ and ASC^-/-^ mice following oral challenge of 10^10^ CFU *E. coli* JM83 12 hours prior to partial hepatectomy. Data shown are the arithmetic mean for the linear scale and geometric mean for the logarithmic scale. Normalized values were analyzed by Student’s t test to compare two experimental groups or by ANOVA to compare more than two groups in parallel. (a-f) Data are representative of 2 independent experiments (4-15 mice per group). (g) Data pooled from 4 independent experiments. The *p*-values are indicated as follows: \**p*≤0.05; ** *p*≤0.01; *** *p*≤0.001; **** *p*≤0.0001.

To investigate the functional relevance of Rorγt^+^ ILC3s in the liver in the systemic protection from intestinally translocated bacteria, we specifically depleted hepatic ILC3s by using anti-Thy1.2 antibody at concentrations that did not affect intestinal Rorγt^+^ ILC3s or NK cells (Fig. 4b)(*10*). After depletion of hepatic Rorγt^+^ ILC3s, translocated *E. coli* JM83 were detected in the livers of SPF C57BL/6 wild-type mice that underwent partial hepatectomy (Fig. 4c), accompanied by a significant decrease in the expression of antimicrobial peptide-encoding genes, suggesting that hepatic Rorγt^+^ ILC3s may be critical for bacterial clearance via the production of antimicrobial peptides (Fig. 4d). The increase in hepatic Rorγt^+^ ILC3s following partial hepatectomy (see Fig. 2e) at 24 hours was preceded by an increase in CD11c^+^ dendritic cells at 12 hours in SPF, but not in germ-free wild-type mice (Fig. 4e and Fig. S12a). We also observed an increase in the gene expression of effector cytokines IL-1β and IL-18 in SPF but not in germ-free wild-type mice subjected to partial hepatectomy (Fig. 4f); these may be secreted by dendritic cells to stimulate Rorγt^+^ ILC3s in the liver in response to intestinal microbes (Fig. 4f, Fig. S12c)(*35, 36*). In order to ascertain the role of IL-1β and IL-18 in bacterial translocation to the liver post surgery, we inoculated *E. coli* JM83 into wild-type mice, IL-18^-/-^ mice, and ASC^-/-^ mice that lack both IL-18 and IL-1β prior to partial hepatectomy. Bacteria were detected in the livers of ASC^-/-^ mice, but not in wild-type or IL-18^-/-^ mice, suggesting that the inflammasome control bacterial translocation (Fig. 4g).

### The microbial-dependent education of the innate immune compartment is required for proper hepatic recovery

Despite the unfavorable ability of intestinal microbes to enter the systemic circulation and cause infections, bacteria are also critical for the development of the host immune system(*20*). It remains unclear to what extent commensal microbiota shape the host healing responses after major surgery(*37*). Partial hepatectomy was therefore performed in germ-free mice, endowed with immature host immune development(*38*). In germ-free wild-type mice, hepatic regeneration was impaired compared to that in SPF wild-type mice, as seen by decreased hepatocellular proliferation (Fig. 5a), suggesting that conditioning of the host by bacteria is critical for optimal outcomes after surgery. Further, also in germ-free mice liver regeneration in *Rag2*^*-/-*^*Il2rγ*^*-/-*^ mice failed in comparison to that in *Rag1*^*-/-*^ mice, suggesting that ILCs play a role in the regenerative process in addition to their role of clearing microbes (Fig. S13a). Comparison of genes upregulated in the liver after partial hepatectomy between germ-free and colonized wild-type mice revealed the activation of specific regenerative pathways such as cell growth and liver differentiation only in colonized wild-type mice, indicating that commensal microbiota are indispensable for triggering specific regenerative responses (Fig. 5b). Since microbiota are known to shape host immune development by synergizing bacterial colonization of the intestinal lumen with penetration of microbial-derived metabolites into systemic organs(*38*), we decided to examine whether activation of these regenerative pathways also occurred in the presence of soluble sterile factors originating from the microbiota. To this end, we transferred serum from colonized or germ-free mice into germ-free recipient mice prior to partial hepatectomy and assessed the expression of those genes that were seen to be specifically up- or downregulated after partial hepatectomy in colonized mice (Fig. S13b). Serum transfer from colonized mice, but not germ-free mice, was sufficient to increase (*Tgif*) or decrease (*Slc41a1* or *Bcl3)* expression in germ-free receiving serum from colonized mice similar to the changes in born and raised colonized mice upon surgery (Fig. 5c), suggesting that bacterial products are sufficient to trigger regenerative pathways. To confirm whether shaping of the host by commensal microbiota is sufficient to improve surgical outcomes, we performed transient colonization of germ-free mice with a genetically modified auxotrophic *E. coli* K-12 HA107 prior to partial hepatectomy. The HA107 strain allows for host immune maturation followed by a return of the mice to germ-free status (Fig. S13c)(*39*). Following partial hepatectomy that was performed 3 weeks after the return to germ-free status, hepatocellular proliferation (Fig. 5d) and expression of cell cycle regulatory genes (Fig. S13d) was increased in transiently colonized wild-type mice, indicating that microbial education improves the regenerative response. However, increase in hepatic proliferation was not observed in *Rag2*^*-/-*^*Il2rγ*^*-/-*^ transiently colonized mice, suggesting that in the absence of an innate and adaptive immune system, commensal microbes fail to trigger hepatic regeneration pathways (Fig. 5e). Consistent with the reported return of transiently colonized mice to germ-free status(*39*), the metabolic profile post-surgery was equivalent between transient microbe-exposed and axenic mice, while both profiles differed from that of SPF mice, suggesting that transient metabolic exposure is sufficient to prime the host immune system for efficient healing (Fig. 5f, Fig. S14, Table S4).

**Fig. 5.**
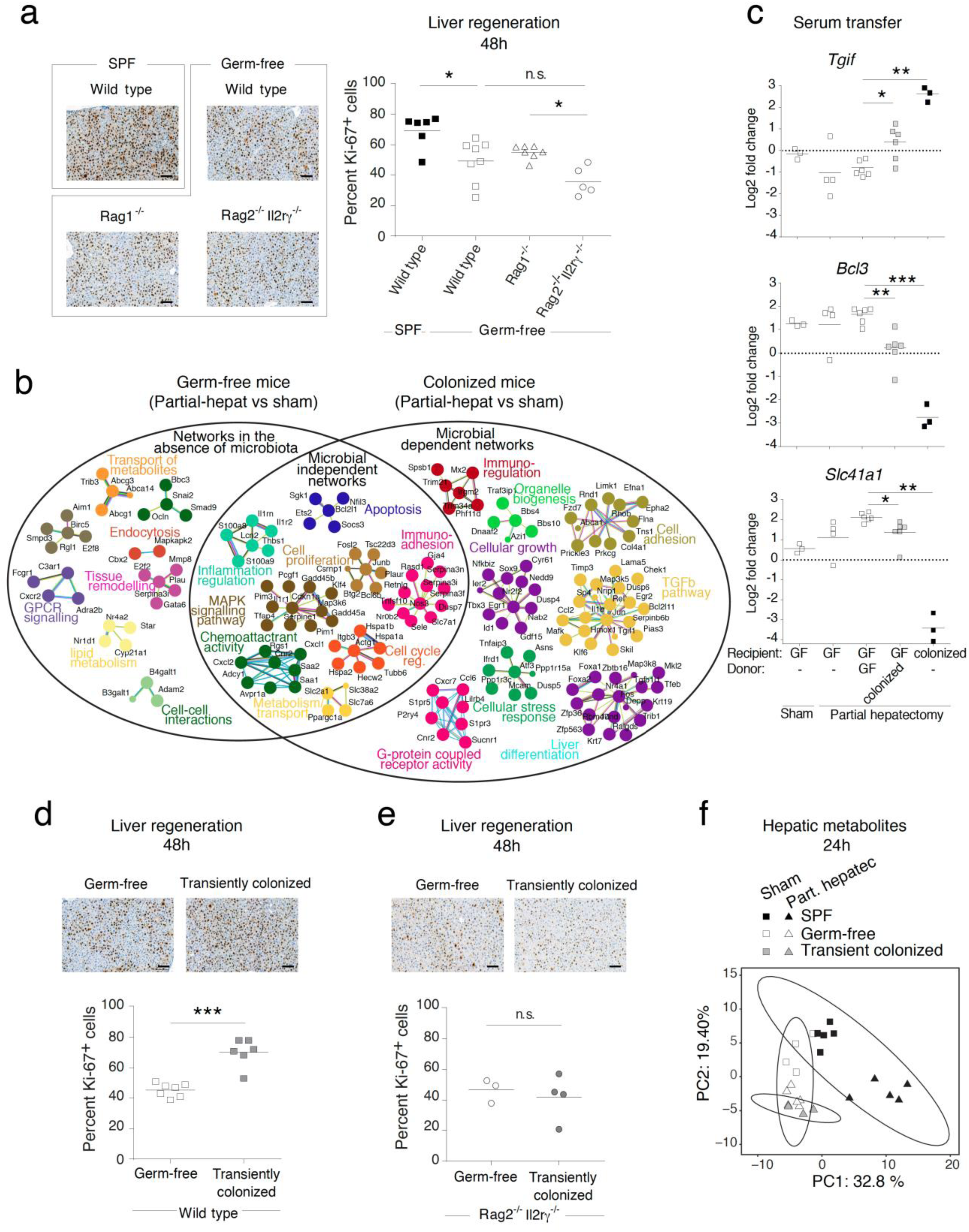
A microbial-dependent education of the innate immune compartments is required for optimal hepatic healing responses. **(a)** SPF wild-type, germ-free wild type, germ-free Rag1^-/-^, and germ-free *Rag2*^*-/-*^ *Il2rγ*^*-/-*^ mice underwent partial hepatectomy and were examined for hepatic regeneration 48 hours after surgery by analyzing the percentage of Ki-67 positive hepatocytes. Representative images are presented on the left and quantification of proliferation is presented on the right. Scale bars in the pictures indicate 100 μm. **(b)** String analysis of upregulated genes in whole liver RNA from germ-free and ASF-colonized mice 3 hours after partial hepatectomy, in comparison to sham-operated mice. **(c)** Sterile serum transfer from germ-free (GF) or colonized donor mice (ASF) into germ-free recipient mice followed by partial hepatectomy and assessment for liver regeneration by analysis of gene expression in whole liver tissue RNA by RT-qPCR and normalized to Gapdh. Sham-operated germ-free mice served as controls. **(d, e)** Germ-free wild-type **(d)** or *Rag2*^*-/-*^*Il2rγ*^*-/-*^ **(e)** mice transiently colonized by four intragastric deliveries of auxotrophic *E. coli* HA107 underwent partial hepatectomy and were analyzed for hepatic regeneration 48 hours after surgery by analyzing the percentage of Ki-67-positive hepatocytes. Representative images are presented on the top and quantification of proliferation is presented at the bottom. Scale bars in the pictures indicate 100 μm **(f)** Principle coordinates analysis of the hepatic metabolome analyzed by high-resolution mass spectrometry in SPF, germ-free and transiently colonized mice 48 hours after partial hepatectomy or sham surgery. The data shown are the arithmetic means for the linear scale. Normalized values were analyzed by Student’s t test to compare two experimental groups or by ANOVA to compare more than two groups in parallel. Data are representative of 2 (a) or 2-3 (b, c) independent experiments (4-15 mice per group). \**p*≤0.05; ** *p*≤0.01; *** *p*≤0.001; n.s.: nonsignificant.

## DISCUSSION

In this paper, we reveal that SSI are not primarily the consequence of contamination during surgery but also a consequence of the failure of intestinal compartmentalization of commensal microbiota (Fig. S15.). We propose that sterile surgery causes an imbalance in intestinal integrity, allowing enteric microbes to translocate and use the host lymphatic system to transit from the intestinal lumen to systemic organs, thereby triggering SSI. Microbial translocation and concomitant SSI are controlled by ILC3s, the inflammasome, and antimicrobial peptides, which together mediate host defenses.

ILC3s have been largely associated with mucosal homeostasis, host-microbe interactions and with the anatomic containment of intestinal microbiota(*12*). For the first time, we show that ILC3s are pivotal for host defense at intestinal mucosal surfaces and systemic defense in organs such as the liver. This study highlights the necessity of resident hepatic ILC3s for systemic bacterial clearance via the induction of antimicrobial peptide production. Thus, in cases of intestinal ILC3 failure, hepatic ILC3s clear penetrated intestinal bacteria, and prevent infections. Because of the lack of antigen-specific receptors, the activation and modulation of ILC function largely depends on signals orchestrated by dendritic cells or macrophages(*40*). Consistent with the function of intestinal ILC3s, we propose that hepatic ILC3 function relies on dendritic cell-dependent inflammasome activation and cytokine signaling to control bacterial clearance (Fig. 4); a process that is potentially similar to IL-22-dependent sensing of bacteria by dendritic cells in the lung(*11*).

Once the intestinal barrier function is disturbed, such as during the acute phase response and associated inflammatory environment, enteric bacteria enter the systemic circulation. Our data show that the small intestine is the main site for bacterial translocation, consistent with prior reports that the barrier function is anatomically and immunologically distinct in different enteric compartments(*28, 29*). The small intestine contains loose mucus and a large surface area to facilitate the absorption of nutrients, and also secretes components such as IgA and antimicrobial peptides to contain enteric microbes within the lumen. These specific properties make the different layers of the small intestine susceptible to acute dysfunction. In contrast, the colon has a dense mucus layer as a consequence of chronic exposure to a large biomass of microbes that makes the colon less sensitive than the intestine to acute challenges(*23, 41*).

We show that Enterobacteriaceae, which are potentially virulent microbial species predominantly found in the intestine(*42*), are the main bacterial family causing SSI irrespective of the site of surgery. These pathobionts are known to cause disease in cases of disturbed intestinal homeostasis, especially in immunocompromised hosts(*43*). We speculate that the failure to adequately keep the bacteria compartmentalized in the lumen, is the cause of pathogenicity rather than dysbiosis or bacterial overgrowth. The extent of surgery, and consequent elevated inflammatory response in the bloodstream, seems to dictate bacterial translocation and eventually infectious complications. *Enterococcus faecalis* caused 10% of the SSIs in our patient cohort and systemically translocated in our mouse model. However, *Enterococcus faecalis* is not displayed in the phylogenetic tree plot because in the 16s rDNA analysis, the frequency was low and the variability was high between volunteers. *Enterococcus faecalis* is highly resistant to antibiotics, forms biofilms and produces highly efficient proteases to resist antimicrobial peptides(*44*). Therefore, despite low abundance in the intestinal biomass bacteria such as intestinal *Enterococcus faecalis* have the ability to trigger SSI.

The finding of intestinal microbiota trafficking via the lymphatic system and thoracic duct into the systemic circulation is supported by a prior clinical report that the accumulation of bacteria in mesenteric lymph nodes after major abdominal surgery was associated with remote SSI(*45*). In addition, the outcome of sepsis has been found to be dependent on gut-derived factors passaging via the lymph and not the venous blood flow(*46*). Interestingly, our data suggest that a large proportion of microbes bypass the splanchnic circulation, making the lung the first filter organ against bacteria. Thus, pneumonia caused by enteric microbes such as *E. coli* may be the consequence of the translocation of microbes originating in the intestine rather than the aspiration of enteric contents. Given that bacteria penetrate via the lymph, antibacterial substances applied with the goal to reduce surgical infections should have high lipophilic availability to be able to reach the lymph in high concentrations in order to control pathogens and prevent the systemic spread of bacteria(*47*).

Current preventive measures against SSI are based on the longstanding dogma that bacterial infections are the result of contamination during surgical procedures. Nevertheless, despite all efforts using hygiene, asepsis, antisepsis and antibiotics, SSI occur in one-tenth of all patients undergoing surgery(*3, 4*). Our studies demonstrate the mechanistic basis for the observation that SSI as well as sepsis have an intestinal microbial signature(*24, 48*). These findings emphasize that the prevention of SSIs requires further understanding of the communication between the intestinal microbiome and systemic host responses. Considering that bacterial translocation and eventual infection are dependent on the immunological status of the host, future studies need to focus on optimizing the immune responses of surgical patients.

Although intestinal microbiota have potentially deleterious effects after translocation and triggering of SSI, they also play key roles in the education of the immune system (*49*). Systemic translocation of intestinal bacteria as indicated by our results, or low amounts of commensal microorganisms as a consequence from standardly prescribed antibiotic prophylaxis prior to surgery, may interfere with recovery. Thus, balanced and optimal interaction of the patient’s immune system with commensal bacteria is critical to achieve successful results after surgery.

Our study addresses SSI, one of the most frequent but under investigated clinical problems and provides mechanistic insights into its pathogenesis. It is highly likely that in a surgical context, compartmentalization of bacteria is ensured by different layers of ILC3s, including the intestinal barrier and, in case of failure, a second line of protection in the liver to clear circulating bacteria. Failure to contain intestinal bacteria by ILC3s and an inflammatory environment may lead to the dissemination of intestinal microbes to remote sites representing a significant cause of infections after surgery. Our study opens new fields of research to reduce surgical infection by targeting patient-specific immune determinants in the intestine and the liver in addition to the efforts to improve measures of hygiene in the surgical environment. Potentially, the SSIs that occur despite all hygiene, antisepsis and asepsis efforts may be reduced by focusing on the host and commensal-dependent factors.

## MATERIAL AND METHODS

### Patient data

Two cohorts of patients were included for the analysis. One dataset includes the patients from a prospective interventional trial, which aimed to test the impact of the introduction of structured communication on SSI as the main outcome parameter (STOP? Trial, NCT02428179). The institutional review board of the Canton of Bern, and Canton Zürich, Switzerland have approved this study (EK ZH 2015-0232, KEK 161/14). The study included all consecutive patients except if patients refused to participate in clinical studies. The outcome parameters SSI was prospectively recorded in the context of a Swiss Surveillance Program (SWISSNOSO) that includes regular monitoring to ensure quality of inclusion of patients and assessment of the endpoints. Data on cultured bacteria were available from 2 participating centers from which the data were included.

All consecutive patients undergoing liver resections for colorectal liver metastasis between 2001 and 2016 at the University hospital of Vienna, Austria, were retrospectively recorded (n=335). The institutional review board of the Medical University of Vienna, Austria has approved this study (424/2010; 2032/2013, NCT01700231, NCT02118545). Infectious complications were summarized with a possible event of an infection during the postoperative course. Major liver resection was defined as resection of three or more liver segments according to the Brisbane 2000 nomenclature (*50*). All bacterial cultures of septic patient were manually reviewed and recorded. Bacterial species were identified by MALDI-TOF mass spectrometry. Other bacteria in fig 1B include: n=2 *Citrobacter spp*, n=1 *Fusobacterium spp*, n=1 *Clostridium sordelli*, n=1 *Streptococcus spp*.

### Mice

Specific-pathogen-free (SPF) wild-type C57BL/6J RccHsd (Envigo), *Rag1*^*-/-*^(*51*) and *Rag2*^*-/-*^*Il2rγ*^*-/-*^(*51, 52*), animals were bred and housed in individual ventilated cages in the Animal Facility of the University of Bern (Switzerland). All experiments were performed in the morning on 8- to 12-week-old adult female mice, supplied with a 12 hour light/dark cycle at 22°C, and fed ad libitum with chow and water.

Altered-Schaedler flora (ASF) *IGIF*^*-/-*^ *(IL-18*^*-/-*^*) and Pycard*^*-/-*^ *(ASC*^*-/-*^) mice were bred and maintained in the same flexible film isolator for more than 10 generations at the Clean Mouse Facility of the University of Bern, Switzerland. The *IGIF*^*-/-*^ *(IL-18*^*-/-*^) was kindly provided by Kevin Maloy (University of Oxford, UK). *Pycard*^*-/-*^ *(ASC*^*-/-*^) were kindly provided by Prof. N. Fasel (University of Lausanne, Department of Biochemistry, Institute for Arthritis Research, Switzerland)(*53*).

Germ-free wild-type C57BL/6J, Rag1^-/-^(*51*) and *Rag2*^*-/-*^*Il2rγ*^*-/-*^(*51, 52*) mice were rederived by axenic two-cell embryo transfer as previously described(*54*) and bred and maintained in flexible film isolators at the Clean Mouse Facility of the University of Bern, Switzerland. Germ-free status was routinely confirmed by extensive aerobic and anaerobic culture as well as DNA stain using SYTOX green (Invitrogen, Gaithersburg, MD) and Gram staining (Harleco, EMD Millipore Corporation, Billerica, MA) of caecal contents to detect unculturable contamination.

All animal procedures were carried out in accordance to the Swiss guidelines for the care and use of laboratory animals, and the experimental protocol received approval by the Animal Care Committee of the Canton of Bern, Switzerland.

### Partial hepatectomy

All surgical procedures were performed under laminar flow and under sterile conditions using general anesthesia with isoflurane (Nicholas Piramal (I) Limited, London, UK). The details of the partial hepatectomy model in mice were previously described(*55*). Briefly, anesthetized mice were immobilized in a supine position and the abdomen was entered through a midline incision. After exposure of liver lobes, partial hepatectomy was performed by central ligature (Vicryl 4-0, Ethicon, Johnson & Johnson, Spreitenbach, Switzerland) of the median and left lobe in order to achieve a standard two-third hepatectomy. The ligated liver lobes were surgically removed, weighted, and further processed. The laparotomy was then closed with a two-layer running suture (Prolene 6-0, Ethicon, Johnson & Johnson, Spreitenbach, Switzerland). During the procedure, the intestine was rinsed with saline to avoid drying-out. Resuscitation of the intraoperative fluid loss was achieved by putting saline into the abdominal cavity at the end of the operation. Analgesia with buprenorphine (Reckitt Benckiser AG, Switzerland) was administered at the beginning of the surgical intervention as well as ad libitum during the postoperative course by subcutaneous injection. At the time of sacrifice, mice were anesthetized by isoflurane inhalation, lethal blood samples were then taken from the inferior vena cava and livers were surgically removed for further analyses. In sham operated animals, a midline incision was performed, the liver was mobilized, the abdominal cavity was rinsed and the abdomen was closed with a two-layer running suture.

### Assessment of bacterial translocation

To determine the presence of bacteria in the liver, lung, spleen, and mesenteric lymph nodes (MLN), the respective organs were harvested under sterile conditions and plated on LB, LB-enriched with streptomycin and BHI (brain-heart infusion)-hemin-protein K-blood agar plates as indicated. LB agar plates were cultured for 24 hours at 37 °C. Blood agar plates were cultured under anaerobic conditions for 48 hours at 37 °C. Bacteria from single isolated colonies were identified by full 16S rRNA sequencing (Sanger sequencing).

### Histological and immunohistochemical analysis of the liver

Immunohistochemistry for hepatocyte proliferation using Ki-67 stain was performed on paraffin-embedded liver sections (3 µm). Fresh liver tissue was fixed with 4% Formaldehyde. Paraffin-embedded tissue sections were then dried, deparaffinized, and rehydrated, followed by blocking of endogenous peroxidase with 3% H2O2 (Sigma H-1009; Sigma-Aldrich Chemie GmbH, Germany) in PBS. Antigen was retrieved by heating the slides for 10 min. Diluted monoclonal mouse anti-Ki-67 antibody (TEC-3 antibody [Dako Code M7249]; Dako, Glostrup, Denmark) was then applied and slides incubated overnight at room temperature in a humidified chamber. Then, biotinylated rabbit anti-mouse IgG ([Code E0464]; Dako, Glostrup, Denmark) was added, followed by a brief incubation with 3,3’-diaminobenzidine substrate (DAB+; [Code K3467]; Dako, Glostrup, Denmark). The tissue sections were counterstained in hematoxylin. Total numbers of Ki-67-labeled hepatocytes were finally determined by counting positively stained hepatocyte nuclei of the whole stained slides in a blinded manner by an independent observer. Percentage of positive proliferating cells (Ki-67^+^ cells) was finally calculated for each animal.

### Bacterial culture

*Escherichia coli* K-12 strain JM83 (F-Δ *(lac-proAB) phi80* Δ *(lacZ)M15 ara rpsL thi* lambda-; streptomycin-resistant)(*56*) was cultured overnight in Luria Broth (LB) medium at 37°C, shaking at 200 rpm. To prepare bacterial inocula for gavage or intravenous injection, bacterial cultures were centrifuged for 10 min at 4000 g and washed twice with sterile PBS. The required dose was resuspended in 500 μl of sterile PBS. JM83-derived auxotrophic *E. coli* K-12 mutant strain HA107 (Δ*asd* Δ*alr* Δ*dadX*)(*39*) was cultured overnight in LB medium containing 100 μg/ml meso-DAP and 400 μg/ml D-alanine at 37°C, 200 rpm.

### Bacterial challenge

Intravenous challenge was performed 6 and 2 days prior to partial hepatectomy. 10^7^ CFU live or heat-killed (70°C, 15 minutes) *E. coli* JM83 were intravenously injected in the tail vein. The heat-killed bacterial inocula were confirmed to be inactivated by culture of aliquots prior to injection on appropriate media. An additional injection of live or heat-killed bacteria was performed 12 hours after partial hepatectomy.

Intraluminal challenge in germ-free or SPF mice was performed with 10^10^ CFU of *E. coli* JM83 by intragastric gavage or by colonic application of the bacteria. The gavage needle was rectally introduced after dilatation of the colon by air to not harm the colonic mucosa and forwarded until the cecum. Partial hepatectomy was performed 12 hours after challenge and mice were sacrificed 24 hours post-surgery.

Transient colonization of adult germ-free mice was performed by administering 4 doses of 10^10^ CFU of *E. coli* HA107 by gavage 3 weeks prior to partial hepatectomy (*39*). After transient colonization, mice were checked to be again germ-free by liquid amplification cultures and microscopic analysis of the feces (as described above).

### Metabolomics analysis and data processing using high-resolution mass spectrometry

Tissues were collected and flash frozen in liquid nitrogen. For extraction of metabolites, 20 volumes/weight cold methanol/acetonitrile/water 2:2:1 (v/v) was added to the samples. Homogenization was performed for 5 min at 25 Hz using stainless steel beads in a tissue homogenizer (Retsch MM 400). Samples were centrifuged at 1000 g for 20 min at 4 °C. The supernatant was collected, transferred in 96-well plate, diluted 1:2 using water and stored at −80 °C until analysis. A pooled quality control (QC) sample was generated by mixing 20 µL of individual tissue sample extracts. The liver extracts were analyzed in randomized order on a 2D UPLC Acquity I-Class system coupled to a quadrupole time-of-flight mass spectrometer (Synapt G2-S HDMS, Waters) operated in positive electrospray ionization (ESI) mode as previously described(*57*). The chromatographic gradient used was slightly adapted: 0 min: 100% mobile phase A at 0.15 mL/min, 1 min: 100% A, 12 min: 1% A, 12.5 min: 1% A at 0.23 mL/min, 13.0 min: 1% A, 13.1 min: 100% A, 13.5 min: 100% A at 0.15 mL/min, 15 min: 100% A. Chromatographic alignment, ion pattern deconvolution (0.45-12 min) and processing including filtering was performed using Progenesis QI software (version 2.2, Nonlinear Dynamics). The metabolic features were searched against the Human Metabolome Database (HMDB, version 4.0) for putative annotation with a mass accuracy of 8 ppm and against an in-house database containing retention times in addition to exact masses. Metabolic features with potential annotations were manually reviewed and features without annotation were excluded from further analysis in the samples comparing germ-free and SPF and HA107 livers. Serum metabolites were extracted as previously described. The liquid chromatography mobile phase was based on acetonitrile instead of methanol and raw data of ESI positive mode were LOESS corrected. Further processing steps were performed similar to the liver tissue extracts. Pathway enriched patterns computing the probability for each pathway was performed using Mummichog(*58*). Compounds with average intensity below 250 in the conditions of interest were removed for the analysis of hepatic metabolites. For Principal Component Analysis of metabolomics data the intensity matrix was log transformed f(x)=log(1+x). The Principal Components were computed with the R function prcomp(*59*). The first two components were displayed with ggplot2(*60*). To display the heatmap metabolomics intensity matrix was normalised f(x_i) = (x_i-min(x))/(max(x)-min(x)), where min(x) denotes the minimum value of a compound over conditions, and max(x) the maximum. The heatmap was displayed with the function heatmap.2 from the R-package gplots using hiearchical cluster for rows with Pearson correlation distance and complete linkage. For the pathway analysis the intensity matrix was log transformed f(x) = log(1+x) and the conditions were compared with student t-test. p-values were corrected by False Discovery Rate. The pathway analysis was done with Mummichog(*58*) using the mass/charge ratio, retention time, p value and log2 fold-change of the transformed data. For the Mummichog analysis of serum samples sham vs partial hepatectomy the pathways of glycosphingolipid and leukotriene metabolism were excluded because low probability of detecting low abundant signalling lipids using this metabolomics approach.

### Cellular isolation

The livers were flushed by injection of cold PBS into the portal vein. Thereafter, the livers were harvested, cut into small pieces with a scalpel and collected for 30 minutes with shaking at 37°C. The medium was then filtered through a 100µm gauge cell strainer into a 50 ml conical tube. Cells were resuspended with cold wash buffer [1X PBS (GIBCO, Life Technologies), 2% Hepes (Sigma Aldrich) and 2% fetal calf serum (FCS)]. Samples were centrifuged at 1250g for 10mins at room temperature. The supernatant was discarded and centrifuged until it became clear. Isolation of cells was achieved by gradient centrifugation using Percoll (GE Healthcare). The cell suspension was resuspended in 40% Percoll (Sigma) solution and layered on top of an 80% Percoll solution. Gradient centrifugation was carried out (2000 g, 20 min, 20°C, no brake). Lymphocytes were collected from the interphase, washed with IMDM (10% FCS) and centrifuged (600 g, 7 min). To isolate leukocytes from the *lamina propria* of small intestine and colon, intestines were removed from the mouse and placed in ice-cold PBS. Residual fat and Peyer’s patches were also removed. The intestine was opened longitudinally and sectioned into 2 cm segments. The tissue was washed once in ice-cold PBS followed by four washes of 8 min in 15 ml of PBS (5 mM EDTA, 10 mM HEPES) with shaking at 37°C to detach epithelial cells. Residual tissue was then washed for 8 min in 15 ml of IMDM containing 10% FCS at 37°C before being minced and digested in 15 ml of IMDM containing 1 mg/ml collagenase type VIII (Sigma) and 10 U/ml DNase I (Roche) with shaking at 37°C for 20-30 min (small intestine) or 30-40 min (colon). The resulting cell suspension was passed through a cell strainer (100 μm) and washed with 20 ml of IMDM 10% FCS. Cells were centrifuged (600 g, 7 min, 4°C) and resuspended in 300 μl of FACS buffer (PBS 2% FCS 2mM EDTA 0.01% NaN3). Intestines were processed individually. Spleens and lymph nodes were cut into small pieces and digested in IMDM (2% FCS) containing collagenase type IA (1 mg/ml, Sigma) and DNase I (10 U/ml, Roche) at 37°C for 30 min. Cellular suspensions were also passed through a cell strainer (40 μm) and washed with IMDM (2% FCS, 2mM EDTA). Cells were finally resuspended in FACS buffer and counted using a Neubauer counting chamber in Trypan blue.

### Cytokines

Lymphocytes were isolated as described earlier. Thereafter, protein isolation was performed using RIPA buffer 10mM Tris, pH8, 1mM EDTA, pH8, 150mM NaCl; phosphatase inhibitors (Na3VO4, NaF, PMSF); a protease inhibitor mix (Sigma-Aldrich) and 0.5% NP40. All extracts were kept on ice for 15 min, centrifuged and the supernatants collected. The total concentration of proteins was quantified with Bradford ultramicroplate reader. After protein quantification, 40 mg of extracted Protein were used for cytokine analysis using U-Plex Mesoscale Cytokine isolation kit from MSD.

### Flow Cytometry

After isolation, cells were washed once with PBS before being stained with fixable viability dye (eBioscience) diluted in DPBS for 30 min on ice. Single cell suspensions were sequentially incubated with primary/biotin- and fluorescence-coupled antibodies diluted in FACS buffer. Cells surface staining was performed by incubating them for 20 minutes at 4°C. Intranuclear staining was performed using the Foxp3 staining kit (eBioscience). The following mouse-specific conjugated antibodies were used: CD127-FITC (A7R34, eBioscience), CD19-BV421, BV785 (6D5, Biolegend), CD3-BV650, BV785 (17A2, Biolegend), CD4-BV785 (RM4-5, Biolegend), CD62L BV510 (MEL-14), Foxp3-AF700 (FJK-16s, eBioscience), Gata3-A647 (TWAJ, BD), NKp46-Percp-Cy55 (19A1.4, Biolegend), RORγt-PE (B2D, eBioscience), T-bet-PECy7 (4B10, eBioscience), fixable viability dye efluor 450 or efluor 506 (ebioscience), CD44-APC Cy7 (Im7, biolegend), CX3CR1-FITC (SA011F11, Biolegend), Siglec F-PE (E50-2440, Biolegend) CD103 Biotinylate (M290, BD), CD11b-APC-Cy7 (M1/70, Biolegend), CD11c PE-CF594 (HL3, BD),F4/80-APC (BM8, Biolegend), MHC-II I-A/I-E AF700 (M5/114.15.2, Biolegend), Ly6C BV421 (HK1.4, eBioscience), Ly6G BV650 (1A8, eBioscience), Streptavin PE Cy7 (BD). LSR Fortessa (BD) was used for flow cytometry and data were analyzed with FlowJo software (Treestar Data Analysis Software).

### RNA isolation

Tissues were removed and snap-frozen using liquid nitrogen. Thawed tissues were immediately homogenized (Retsch bead-beater) in 500 μl of Nucleozol reagent. Chloroform (200 μl) was added, samples were mixed, and centrifuged (12,000 g, 15 min, 4°C). The upper phase was collected and RNA was precipitated with ice-cold isopropanol by centrifugation (12,000 g, 10 min, 4°C). The RNA pellet was washed with 75% ethanol, dried and resuspended in RNase-free water. RNA concentrations and purity were analyzed using a Nanodrop2000 (Thermo Scientific).

### RT-qPCR

Genomic DNA in RNA samples was digested using the DNA-free kit (Ambion) and RNA was reverse-transcribed into cDNA using Superscript III reverse transcriptase according to Invitrogen protocols. Real-time PCR was performed using 90 well-plates with 50-100 ng of cDNA per well containing the SsoFast EvaGreen Supermix (Biorad) and gene-specific primers. All reactions were run in triplicate. Samples were normalized to the expression of TBP for TaqMan and GAPDH for SYBR Green by calculating 2^ (-deltaCt).

#### Primer sequences

*GAPDH* (Gapdh) forward: 5‘ - CAT CAA GAA GGT GGT GAA GC - 3‘

*GAPDH* (Gapdh) reverse: 5‘ - CCT GTT GCT GTA GCC GTA TT - 3‘

*REG3B* (Reg3beta) forward: 5’ – GCA GAA CCC AAT GGA GGT GG - 3’

*REG3B* (Reg3beta) reverse: 5’ – CAC CCA GGG ATG TGA GAA GAG - 3’

*REG3G* (Reg3gamma) forward: 5’ – TTC CTG TCC TCC ATG ATC AAA - 3’

*REG3G* (Reg3gamma) reverse: 5’ – CAT CCA CCT CTG TTG GGT TC - 3’

*LYZ1* (Lysozyme 1) forward: 5’ – CTT GTC ACT CCT CAC CCC TG - 3’

*LYZ1* (Lysozyme 1) reverse: 5’ – AGC CGT TCC CCT TCC AAT G - 3’

*Il22* (IL-22) forward: 5’ – ATG AGT TTT TCC CTT ATG GGG AC- 3’

*Il22* (IL-22) reverse: 5’ – GCT GGA AGT TGG ACA CCT CAA- 3’

*DEFA21* (alpha defensin 21) forward: 5’ – AGG CTG TGT CTG TCT CCT TTG - 3’

*DEFA21* (alpha defensin 21) reverse: 5’ – TGC AAG CAT CCA TCA CAC TGG - 3’

*Il6 (IL-6)* forward: 5’ - TCG GAG GCT TAA TTA CAC ATG TTC T - 3’

*Il6 (IL-6)* reverse: 5’ - GCA TCA TCG TTG TTC ATA CAA TCA - 3’

*Il1B* (IL-1β, forward: 5’ - ACCTGTCCTGTGTAATGAAAGACG - 3’

*Il1B* (IL-1β) reverse: 5’ - TGGGTATTGCTTGGGATCCA - 3’

*Il18* (IL-18) forward: 5’ - ACA ACT TTG GCC GAC TTC AC - 3’

*Il18* (IL-18) reverse: 5’ - TGG ATC CAT TTC CTC AAA GG - 3’

*CLDN1* (Claudin 1) forward: 5’ - ACT CCT TGC TGA ATC TGA ACA GT - 3’

*CLDN1* (Claudin 1) reverse: 5’ - GGA CAC AAA GAT TGC GAT CAG - 3’

*CLDN8* (Claudin 8) forward: 5’ - GGA GGA CGA TGG CAA CCT AC - 3’

*CLDN8* (Claudin 8) reverse: 5’ - AGC GGT TCT CAA ACA CCA CA - 3’

*CLDN15* (Claudin 15) forward: 5’ - AAC GTG GGC AAC ATG GAT CT - 3’

*CLDN15* (Claudin 15) reverse: 5’ - CGT ACC ACG AGA TAG CCA CC - 3’

*TJP1* (Zo1) forward: 5’ - AGA CGC CCG AGG GTG TAG - 3’

*TJP1* (Zo1) reverse: 5’ - TGG GAG AAA AGT CCG GGA AG - 3’

*TJP3* (Zo3) forward: 5’ - CAG GTC GAT CAT GGG GTG AG - 3’

*TJP3* (Zo3) reverse: 5’ - CCA GAC CGT TGG CTT CAG AT - 3’

For the assessment of hepatocyte proliferation, the quantitative RT–PCR with TaqMan gene expression assays were used. Reagents were used according to the standard protocols (Applied Biosystem). Mouse probes used include *BIRC5* (apoptosis inhibitor 4) (Mm00599749_m1), *CDKN1A* (cyclin dependent kinase inhibitor 1A) (Mm004324480), *CCNA2* (cyclin A2) (Mm00438063_m1), *CCND1* (cyclin D1) (Mm00432359), *FOXM1B* (Mm00514925_m1), and *TBP* (Mm01277042_m1).

### Microbial community analysis

The mucosa-associated microbiota of fecal and skin samples from two different family was analyzed by 16S rRNA amplicon sequencing on the lonTorrent PGM™ platform, as previously described(*61*). Briefly, microbial DNA was extracted from fecal and skin samples using the QIAamp DNA stool kit (Qiagen) according to the manufacturer’s instructions(*41*). Concentrations of purified DNA were measured using NanoDrop (ThermoScientific). DNA was pooled at a concentration of 26pM and was sequenced for the V5/V6 region of 16S rRNA genes using a multiplex approach with the barcoded forward fusion primer 5′-*CCATCTCATCCCTGCGTGTCTCCGACTCAG* BARCODE ATTAGATACCCYGGTAGTCC-3′ in combination with the reverse fusion primer 5′-*CCTCTCTATGGGCAGTCGGTGAT*ACG AGCTGACGACARCCATG-3′, in IonTorrent PGM system according to the manufacturer’s instructions (ThermoFisher)(*62, 63*). Data was further analyzed using QIIME *v1.9.1* pipeline after filtering out low quality (accuracy of base calling; q<25) samples. Operational taxonomic units (OTUs) were picked using UCLUST with a 97% sequence identity threshold followed by taxonomy assignment using the latest GreenGenes database from May 2013 (http://greengenes.secondgenome.com/downloads). Alpha diversity (that describes the number of different taxa within a sample) and beta diversity (that delineates differences between samples (i.e. “between habitat” diversity) were calculated using the *phyloseq* pipeline in R (*v3.4*)(*64, 65*). The non-parametric Mann-Whitney U-tests was used to compare alpha diversity between samples and Adonis from vegan package to assess the effects of groups for beta diversity via *phyloseq* in R(*64, 65*). Taxonomic differences at phylum and genus levels between tested groups were identified using the “multivariate analysis by linear models” (MaAsLin) R package(*66*). Only taxa present in at least 50% of samples and OTUs comprising more than 0.001% of relative abundance were analyzed in MaAsLin. A p< 0.05 and a false discovery rate (FDR; Benjamini-Hochberg correction) of q < 0.05 were set as cut-off values for significance. All the relevant codes for running the MaAsLin in R platform are available in Dr. Huttenhower’s group webpage (https://huttenhower.sph.harvard.edu). Phylogenetic tree was plot using GraPhlAn(*67*).

### Confocal endomicroscopy (Cellvizio®) experiments

Partial hepatectomy was performed immediately prior to imaging. After the operation, 70 kD Fluorescein isothiovyanate (FITC)-dextran (10 ug/ml, Sigma) in PBS was intravenously injected into the tail vein to provide real-time contrast. Immediately after injection, the terminal ileum was incised and a confocal miniprope (Cellvizio, Mauna Kea Technologies, Paris, France) was introduced into the small intestine. The fluorescence intensity outside:inside was determined using ImageJ software (NIH) calculating the ratios of extravascular to intravascular fluorescence.

### Fecal albumin intestinal permeability assays

Fecal pellets were homogenized in sterile PBS measured by ELISA following the protocol of a commercially available kit (Bethyl).

### Thoracic duct ligation

Thoracic duct ligation was performed as previously described with some alterations(*68*). After gavage of 200 ul milk cream (25% fat) 30 minutes prior to the operation, the anesthesized mice were placed on the right side and a left-sided subcostal incision was performed. After placement of a self-retaining retractor, the aorta was exposed. After incision of the dorsal retroperitoneum, the aorta was freed from surrounding fatty tissue. The thoracic duct was exposed and carefully freed from the aorta. A metal clip was then placed for closure of the thoracic duct (Clip 9 Vitalitec, Peters Surgical, Bobigny Cedex, France). The incision was closed with a two-layer running suture.

### Thy1.2 depletion

Anti-CD90.2 mAb (Thy1.2, 30H12) was purchased from BioXCell (West Lebanon, NH). Depletion mAb or isotype mAb (rat anti-mouse IgG) treatments were administered i.p. 6 days, 3 days and 1 day prior to the surgery at a dose of 250 µg/mouse(*10*).

### RNA sequencing and network analysis

RNA was isolated from the liver collected 3 hours after sham and partial hepatectomy of the germ-free or colonized mice, and was purified using the RNeasy MinElute kit (Qiagen). The RNA concentration and quality was determined using a Bioanalyzer 2100 (Agilent). Library preparation was performed using TruSeq RNA sample preparation v2 kit (TruSeq Stranded mRNA Sample Preparation, Illumina). Libraries were sequenced by Illumina HiSeq 2500 on the 100 bp paired-end mode. The quality and number of reads were documented and all reads were mapped to the mouse genome using tophat2. Htseq_count software was used to evaluate the gene-specific reads (http://www.ensembl.org/index.html). These hits per gene were compared across the different experimental groups using the DESeq2 package in R. Adjusted log2(fold-change) and p values (corrected for multiple tests)(*69*) were used to determine significant differences in gene expression. For each RNASeq dataset, genes were scored according to an absolute fold-change ≥ 2 and adjusted p-value of < 0.01. The proteins corresponding to the obtained gene sets were searched against the version 10 of the STRING database(*70*) to display functional protein-association networks. Interactions were considered with a STRING confidence score ≥ 0.4 (medium 9 and high confidence). Multiple rounds of iteration of the k-means clustering method were performed.

### Serum transfer

Serum was collected under sterile conditions from germ-free or colonized mice. Serum was sterile-filtered and transferred into germ-free recipients 7 days and 3 days before, and at the day of the surgery (3 x 250 μl per mouse) by intravenous injection into the tail vein.

### Statistical analysis

Dot plots with a logarithmic scale show the geometric mean of each experimental group in addition to the individual samples represented as single data points. Dot plots with a linear scale show the arithmetic mean. Normalized values were analyzed by Student’s t test or one-way ANOVA. P-values are indicated as follows: * p ≤ 0.05; ** p ≤ 0.01; *** p ≤ 0.001, **** p ≤ 0.0001.

## SUPPLEMENTARY MATERIALS

**Fig. S1.** Gating strategy for flow cytometry analysis of innate lymphoid cells/Th17 cells.

**Fig. S2.** Minor changes in other ILC population in the small intestine, colon, MLN and liver

**Fig. S3.** Major liver surgery alters ILC3s only in the presence of microbiota

**Fig. S4.** Systemic bacteria originate from the intestine in the context of major liver surgery

**Fig. S5.** Circulating intestinal bacteria impair liver regeneration

**Fig. S6.** Systemic dissemination of intestinal bacteria is independent of the microbial composition

**Fig. S7.** Pathway analysis of serum metabolic changes after partial hepatectomy

**Fig. S8.** Bacterial challenge does not impair liver regeneration in SPF wild type mice

**Fig. S9.** Enhanced induction of tight junction mRNA expression in addition to a cytokine immune response in the colon in comparison to the small intestine

**Fig. S10.** Hepatic metabolites do not follow the trajectory of bacteria

**Fig. S11.** Expression of antimicrobial peptides in the small intestine and colon are not affected upon partial hepatectomy

**Fig. S12.** Presence of translocated bacteria induced by the surgery enhance inflammatory cytokine secretion in the liver

**Fig. S13.** Absence of intestinal commensal microbiota decrease liver regeneration

**Fig. S14.** Distinct differences in metabolic pathways between germ-free and colonized mice

**Fig. S15.** Graphical abstract

**Table S1.** Raw data - Serum metabolome 8 hours post-surgery in colonized wild type mice

**Table S2.** Raw data - Hepatic metabolome 24 hours post-surgery in wild type, and *Rag2*^*-/-*^*Il2rγ*^*-/-*^ mice +/- thoracic duct ligation

**Table S3.** Raw data – Patient database

**Table S4.** Raw data – RNA-Sequencing of whole liver tissue 3 hours post-surgery in colonized and germ-free wild type mice

**Table S5.** Raw data - Hepatic metabolome sham/12hours and 24 hours post-surgery in colonized and germ-free wild type mice

**Movie S1.** Vascular permeability upon major liver surgery in C57BL/6 wild type mice

**Movie S2.** Vascular permeability upon major liver surgery in *Rag2*^*-/-*^*Il2rγ*^*-/-*^

## ACKNOWLEDGMENTS

We thank members of the research laboratory of the visceral surgery and medicine, Dr. S. Ganal-Vonarburg and Prof. A.J. Macpherson for critical input and support. We thank I. Büchi for the technical support. Prof. R. Wiest and Dr. M. Sorribas helped with the Cellvizio experiments. The clean mouse facility of the department of Biomedical Research kindly housed and provided the GF mice. Dr. Francesca Ronchi facilitated the use of *IGIF (IL-18)*^*-/-*^ mice. MOJ received funding by the ARS Bridge to academic career grant. MOJ, GB, MGA and SH receive funding by the Swiss National Science foundation (Grant-ID: MOJ 184425, GB 166594, 156882, MGA 168012, SH 138452, 169791).

## AUTHOR CONTRIBUTIONS

M.O.J., T.M., and M.G.A. performed the animal experiments. M.O.J., T.M., M.G.A. and G. B. made the experimental plan of investigation. M.O.J., M.G.A, G.B. analyzed the experimental data and wrote the manuscript. L.S. did RNA isolation and PCR experiments. B.Y. carried out the microbiota profiling analysis on fecal and skin samples. D.S.T., B.Y. and C.M. performed and interpreted the bioinformatic analysis. D.S. and B.Y. performed the sequencing of human skin and fecal samples. S.H. provided the *E. coli* K-12 HA107. S.H., D.C., D.S. provided input during work discussions and provided guidance on the concept and design. K.F. and M.M. were responsible for the metabolomic analysis and its interpretation. G.B., F.T. conceived, designed and were responsible for the clinical study. P.S. and D.P. provided and interpreted the human data on liver surgeries. M.O.J. and M.G.A. created the figures, wrote the methods.

All authors edited and approved the manuscript.

## DATA AVAILABILITY STATEMENT

The data that support the findings of this study are available from the corresponding author upon reasonable request. Upon publication, the raw data will be published in the Bern Open Repository and Information System (BORIS).

## REFERENCES

1. lancetglobalsurgery.org, Number of surgical procedures (per 100,000 population).

2. S. S. Magill, E. O’Leary, S. J. Janelle, D. L. Thompson, G. Dumyati, J. Nadle, L. E. Wilson, M. A. Kainer, R. Lynfield, S. Greissman, S. M. Ray, Z. Beldavs, C. Gross, W. Bamberg, M. Sievers, C. Concannon, N. Buhr, L. Warnke, M. Maloney, V. Ocampo, J. Brooks, T. Oyewumi, S. Sharmin, K. Richards, J. Rainbow, M. Samper, E. B. Hancock, D. Leaptrot, E. Scalise, F. Badrun, R. Phelps, J. R. Edwards, T. Emerging Infections Program Hospital Prevalence Survey, Changes in Prevalence of Health Care-Associated Infections in U.S. Hospitals. The New England journal of medicine 379, 1732-1744 (2018).

3. B. Allegranzi, B. Zayed, P. Bischoff, N. Z. Kubilay, S. de Jonge, F. de Vries, S. M. Gomes, S. Gans, E. D. Wallert, X. Wu, M. Abbas, M. A. Boermeester, E. P. Dellinger, M. Egger, P. Gastmeier, X. Guirao, J. Ren, D. Pittet, J. S. Solomkin, W. H. O. G. D. Group, New WHO recommendations on intraoperative and postoperative measures for surgical site infection prevention: an evidence-based global perspective. Lancet Infect Dis 16, e288-e303 (2016).

4. K. Reinhart, R. Daniels, N. Kissoon, F. R. Machado, R. D. Schachter, S. Finfer, Recognizing Sepsis as a Global Health Priority - A WHO Resolution. The New England journal of medicine 377, 414-417 (2017).

5. N. Faraday, P. Rock, E. E. Lin, T. M. Perl, K. Carroll, T. Stierer, P. Robarts, A. McFillin, T. Ross, A. S. Shah, L. H. Riley, R. J. Tamargo, J. H. Black, E. Blasco-Colmenares, E. Guallar, Past history of skin infection and risk of surgical site infection after elective surgery. Annals of surgery 257, 150-154 (2013).

6. S. I. Berrios-Torres, C. A. Umscheid, D. W. Bratzler, B. Leas, E. C. Stone, R. R. Kelz, C. E. Reinke, S. Morgan, J. S. Solomkin, J. E. Mazuski, E. P. Dellinger, K. M. F. Itani, E. F. Berbari, J. Segreti, J. Parvizi, J. Blanchard, G. Allen, J. Kluytmans, R. Donlan, W. P. Schecter, C. Healthcare Infection Control Practices Advisory, Centers for Disease Control and Prevention Guideline for the Prevention of Surgical Site Infection, 2017. JAMA surgery 152, 784-791 (2017).

7. J. C. Hernaiz-Leonardo, M. F. Golzarri, P. Cornejo-Juarez, P. Volkow, C. Velazquez, M. Ostrosky-Frid, D. Vilar-Compte, Microbiology of surgical site infections in patients with cancer: A 7-year review. American journal of infection control 45, 761-766 (2017).

8. L. V. Hooper, D. R. Littman, A. J. Macpherson, Interactions between the microbiota and the immune system. Science 336, 1268-1273 (2012).

9. D. Ramanan, K. Cadwell, Intrinsic Defense Mechanisms of the Intestinal Epithelium. Cell host & microbe 19, 434-441 (2016).

10. G. F. Sonnenberg, L. A. Monticelli, T. Alenghat, T. C. Fung, N. A. Hutnick, J. Kunisawa, N. Shibata, S. Grunberg, R. Sinha, A. M. Zahm, M. R. Tardif, T. Sathaliyawala, M. Kubota, D. L. Farber, R. G. Collman, A. Shaked, L. A. Fouser, D. B. Weiner, P. A. Tessier, J. R. Friedman, H. Kiyono, F. D. Bushman, K. M. Chang, D. Artis, Innate lymphoid cells promote anatomical containment of lymphoid-resident commensal bacteria. Science 336, 1321-1325 (2012).

11. J. Gray, K. Oehrle, G. Worthen, T. Alenghat, J. Whitsett, H. Deshmukh, Intestinal commensal bacteria mediate lung mucosal immunity and promote resistance of newborn mice to infection. Science translational medicine 9, (2017).

12. E. Vivier, D. Artis, M. Colonna, A. Diefenbach, J. P. Di Santo, G. Eberl, S. Koyasu, R. M. Locksley, A. N. J. McKenzie, R. E. Mebius, F. Powrie, H. Spits, Innate Lymphoid Cells: 10 Years On. Cell 174, 1054-1066 (2018).

13. D. R. Withers, M. R. Hepworth, Group 3 Innate Lymphoid Cells: Communications Hubs of the Intestinal Immune System. Frontiers in immunology 8, 1298 (2017).

14. S. P. Broadley, A. Plaumann, R. Coletti, C. Lehmann, A. Wanisch, A. Seidlmeier, K. Esser, S. Luo, P. C. Ramer, S. Massberg, D. H. Busch, M. van Lookeren Campagne, A. Verschoor, Dual-Track Clearance of Circulating Bacteria Balances Rapid Restoration of Blood Sterility with Induction of Adaptive Immunity. Cell host & microbe 20, 36-48 (2016).

15. A. J. Macpherson, M. Heikenwalder, S. C. Ganal-Vonarburg, The Liver at the Nexus of Host-Microbial Interactions. Cell host & microbe 20, 561-571 (2016).

16. R. Kudira, T. Malinka, A. Kohler, M. Dosch, M. G. de Aguero, N. Melin, S. Haegele, P. Starlinger, N. Maharjan, S. Saxena, A. Keogh, D. Stroka, D. Candinas, G. Beldi, P2X1-regulated IL-22 secretion by innate lymphoid cells is required for efficient liver regeneration. Hepatology 63, 2004-2017 (2016).

17. M. Forkel, L. Berglin, E. Kekalainen, A. Carlsson, E. Svedin, J. Michaelsson, M. Nagasawa, J. S. Erjefalt, M. Mori, M. Flodstrom-Tullberg, A. Bergquist, H. G. Ljunggren, M. Westgren, U. Lindforss, D. Friberg, C. Jorns, E. Ellis, N. K. Bjorkstrom, J. Mjosberg, Composition and functionality of the intrahepatic innate lymphoid cell-compartment in human nonfibrotic and fibrotic livers. Eur J Immunol 47, 1280-1294 (2017).

18. A. Matsumoto, T. Kanai, Y. Mikami, P. S. Chu, N. Nakamoto, H. Ebinuma, H. Saito, T. Sato, H. Yagita, T. Hibi, IL-22-producing RORgammat-dependent innate lymphoid cells play a novel protective role in murine acute hepatitis. PloS one 8, e62853 (2013).

19. Q. Wang, J. Zhou, B. Zhang, Z. Tian, J. Tang, Y. Zheng, Z. Huang, Y. Tian, Z. Jia, Y. Tang, J. C. van Velkinburgh, Q. Mao, X. Bian, Y. Ping, B. Ni, Y. Wu, Hepatitis B virus induces IL-23 production in antigen presenting cells and causes liver damage via the IL-23/IL-17 axis. PLoS pathogens 9, e1003410 (2013).

20. A. J. Macpherson, M. G. de Aguero, S. C. Ganal-Vonarburg, How nutrition and the maternal microbiota shape the neonatal immune system. Nature reviews. Immunology 17, 508-517 (2017).

21. M. J. Xu, D. Feng, H. Wu, H. Wang, Y. Chan, J. Kolls, N. Borregaard, B. Porse, T. Berger, T. W. Mak, J. B. Cowland, X. Kong, B. Gao, Liver is the major source of elevated serum lipocalin-2 levels after bacterial infection or partial hepatectomy: a critical role for IL-6/STAT3. Hepatology 61, 692-702 (2015).

22. M. A. Krezalek, S. Hyoju, A. Zaborin, E. Okafor, L. Chandrasekar, V. Bindokas, K. Guyton, C. P. Montgomery, R. S. Daum, O. Zaborina, S. Boyle-Vavra, J. C. Alverdy, Can Methicillin-resistant Staphylococcus aureus Silently Travel From the Gut to the Wound and Cause Postoperative Infection? Modeling the “Trojan Horse Hypothesis”. Annals of surgery 267, 749-758 (2018).

23. F. Melo-Gonzalez, H. Kammoun, E. Evren, E. E. Dutton, M. Papadopoulou, B. M. Bradford, C. Tanes, F. Fardus-Reid, J. R. Swann, K. Bittinger, N. A. Mabbott, B. A. Vallance, T. Willinger, D. R. Withers, M. R. Hepworth, Antigen-presenting ILC3 regulate T cell-dependent IgA responses to colonic mucosal bacteria. J Exp Med 216, 728-742 (2019).

24. F. B. Tamburini, T. M. Andermann, E. Tkachenko, F. Senchyna, N. Banaei, A. S. Bhatt, Precision identification of diverse bloodstream pathogens in the gut microbiome. Nature medicine 24, 1809-1814 (2018).

25. Y. T. Xiao, W. H. Yan, Y. Cao, J. K. Yan, W. Cai, Neutralization of IL-6 and TNF-alpha ameliorates intestinal permeability in DSS-induced colitis. Cytokine 83, 189-192 (2016).

26. F. Raimondi, P. Santoro, M. V. Barone, S. Pappacoda, M. L. Barretta, M. Nanayakkara, C. Apicella, L. Capasso, R. Paludetto, Bile acids modulate tight junction structure and barrier function of Caco-2 monolayers via EGFR activation. American journal of physiology. Gastrointestinal and liver physiology 294, G906-913 (2008).

27. I. Spadoni, E. Zagato, A. Bertocchi, R. Paolinelli, E. Hot, A. Di Sabatino, F. Caprioli, L. Bottiglieri, A. Oldani, G. Viale, G. Penna, E. Dejana, M. Rescigno, A gut-vascular barrier controls the systemic dissemination of bacteria. Science 350, 830-834 (2015).

28. S. A. Houston, V. Cerovic, C. Thomson, J. Brewer, A. M. Mowat, S. Milling, The lymph nodes draining the small intestine and colon are anatomically separate and immunologically distinct. Mucosal immunology 9, 468-478 (2016).

29. D. Esterhazy, M. C. C. Canesso, L. Mesin, P. A. Muller, T. B. R. de Castro, A. Lockhart, M. ElJalby, A. M. C. Faria, D. Mucida, Compartmentalized gut lymph node drainage dictates adaptive immune responses. Nature 569, 126-130 (2019).

30. M. L. Balmer, E. Slack, A. de Gottardi, M. A. Lawson, S. Hapfelmeier, L. Miele, A. Grieco, H. Van Vlierberghe, R. Fahrner, N. Patuto, C. Bernsmeier, F. Ronchi, M. Wyss, D. Stroka, N. Dickgreber, M. H. Heim, K. D. McCoy, A. J. Macpherson, The liver may act as a firewall mediating mutualism between the host and its gut commensal microbiota. Science translational medicine 6, 237ra266 (2014).

31. H. X. Liu, C. S. Rocha, S. Dandekar, Y. J. Wan, Functional analysis of the relationship between intestinal microbiota and the expression of hepatic genes and pathways during the course of liver regeneration. Journal of hepatology 64, 641-650 (2016).

32. W. Xiao, M. Ren, C. Zhang, S. Li, W. An, Amelioration of nonalcoholic fatty liver disease by hepatic stimulator substance via preservation of carnitine palmitoyl transferase-1 activity. American journal of physiology. Cell physiology 309, C215-227 (2015).

33. C. Xie, S. Takahashi, C. N. Brocker, S. He, C. Li, G. Xie, K. Jang, X. Gao, K. W. Krausz, A. Qu, M. Levi, F. J. Gonzalez, Hepatocyte peroxisome proliferator-activated receptor alpha regulates bile acid synthesis and transport. Biochimica et biophysica acta. Molecular and cell biology of lipids, (2019).

34. L. J. Zhang, R. L. Gallo, Antimicrobial peptides. Current biology: CB 26, R14-19 (2016).

35. A. Geremia, C. V. Arancibia-Carcamo, Innate Lymphoid Cells in Intestinal Inflammation. Frontiers in immunology 8, 1296 (2017).

36. F. Melo-Gonzalez, M. R. Hepworth, Functional and phenotypic heterogeneity of group 3 innate lymphoid cells. Immunology 150, 265-275 (2017).

37. X. Wu, R. Sun, Y. Chen, X. Zheng, L. Bai, Z. Lian, H. Wei, Z. Tian, Oral ampicillin inhibits liver regeneration by breaking hepatic innate immune tolerance normally maintained by gut commensal bacteria. Hepatology 62, 253-264 (2015).

38. Y. Uchimura, T. Fuhrer, H. Li, M. A. Lawson, M. Zimmermann, B. Yilmaz, J. Zindel, F. Ronchi, M. Sorribas, S. Hapfelmeier, S. C. Ganal-Vonarburg, M. Gomez de Aguero, K. D. McCoy, U. Sauer, A. J. Macpherson, Antibodies Set Boundaries Limiting Microbial Metabolite Penetration and the Resultant Mammalian Host Response. Immunity 49, 545-559 e545 (2018).

39. S. Hapfelmeier, M. A. Lawson, E. Slack, J. K. Kirundi, M. Stoel, M. Heikenwalder, J. Cahenzli, Y. Velykoredko, M. L. Balmer, K. Endt, M. B. Geuking, R. Curtiss, 3rd, K. D. McCoy, A. J. Macpherson, Reversible microbial colonization of germ-free mice reveals the dynamics of IgA immune responses. Science 328, 1705-1709 (2010).

40. M. Ebbo, A. Crinier, F. Vely, E. Vivier, Innate lymphoid cells: major players in inflammatory diseases. Nature reviews. Immunology 17, 665-678 (2017).

41. H. Li, J. P. Limenitakis, T. Fuhrer, M. B. Geuking, M. A. Lawson, M. Wyss, S. Brugiroux, I. Keller, J. A. Macpherson, S. Rupp, B. Stolp, J. V. Stein, B. Stecher, U. Sauer, K. D. McCoy, A. J. Macpherson, The outer mucus layer hosts a distinct intestinal microbial niche. Nature communications 6, 8292 (2015).

42. N. Kamada, G. Y. Chen, N. Inohara, G. Nunez, Control of pathogens and pathobionts by the gut microbiota. Nature immunology 14, 685-690 (2013).

43. J. Chow, S. K. Mazmanian, A pathobiont of the microbiota balances host colonization and intestinal inflammation. Cell host & microbe 7, 265-276 (2010).

44. O. Nesuta, M. Budesinsky, R. Hadravova, L. Monincova, J. Humpolickova, V. Cerovsky, How proteases from Enterococcus faecalis contribute to its resistance to short alpha-helical antimicrobial peptides. Pathogens and disease 75, (2017).

45. E. Nishigaki, T. Abe, Y. Yokoyama, M. Fukaya, T. Asahara, K. Nomoto, M. Nagino, The detection of intraoperative bacterial translocation in the mesenteric lymph nodes is useful in predicting patients at high risk for postoperative infectious complications after esophagectomy. Annals of surgery 259, 477-484 (2014).

46. E. A. Deitch, Role of the gut lymphatic system in multiple organ failure. Curr Opin Crit Care 7, 92-98 (2001).

47. A. Bogoslowski, P. Kubes, Lymph Nodes: The Unrecognized Barrier against Pathogens. ACS Infect Dis 4, 1158-1161 (2018).

48. J. Defazio, I. D. Fleming, B. Shakhsheer, O. Zaborina, J. C. Alverdy, The opposing forces of the intestinal microbiome and the emerging pathobiome. Surg Clin North Am 94, 1151-1161 (2014).

49. S. P. Spencer, G. K. Fragiadakis, J. L. Sonnenburg, Pursuing Human-Relevant Gut Microbiota-Immune Interactions. Immunity 51, 225-239 (2019).

50. Y. Y. Pang, The Brisbane 2000 terminology of liver anatomy and resections. HPB 2000; 2:333-39. HPB (Oxford) 4, 99; author reply 99-100 (2002).

51. P. Mombaerts, J. Iacomini, R. S. Johnson, K. Herrup, S. Tonegawa, V. E. Papaioannou, RAG-1-deficient mice have no mature B and T lymphocytes. Cell 68, 869-877 (1992).

52. F. Colucci, C. Soudais, E. Rosmaraki, L. Vanes, V. L. Tybulewicz, J. P. Di Santo, Dissecting NK cell development using a novel alymphoid mouse model: investigating the role of the c-abl proto-oncogene in murine NK cell differentiation. Journal of immunology 162, 2761-2765 (1999).

53. S. K. Drexler, L. Bonsignore, M. Masin, A. Tardivel, R. Jackstadt, H. Hermeking, P. Schneider, O. Gross, J. Tschopp, A. S. Yazdi, Tissue-specific opposing functions of the inflammasome adaptor ASC in the regulation of epithelial skin carcinogenesis. Proceedings of the National Academy of Sciences of the United States of America 109, 18384-18389 (2012).

54. E. Slack, S. Hapfelmeier, B. Stecher, Y. Velykoredko, M. Stoel, M. A. Lawson, M. B. Geuking, B. Beutler, T. F. Tedder, W. D. Hardt, P. Bercik, E. F. Verdu, K. D. McCoy, A. J. Macpherson, Innate and adaptive immunity cooperate flexibly to maintain host-microbiota mutualism. Science 325, 617-620 (2009).

55. D. Inderbitzin, P. Studer, D. Sidler, G. Beldi, V. Djonov, A. Keogh, D. Candinas, Regenerative capacity of individual liver lobes in the microsurgical mouse model. Microsurgery 26, 465-469 (2006).

56. J. Vieira, J. Messing, The pUC plasmids, an M13mp7-derived system for insertion mutagenesis and sequencing with synthetic universal primers. Gene 19, 259-268 (1982).

57. B. Rindlisbacher, C. Schmid, T. Geiser, C. Bovet, M. Funke-Chambour, Serum metabolic profiling identified a distinct metabolic signature in patients with idiopathic pulmonary fibrosis -a potential biomarker role for LysoPC. Respiratory research 19, 7 (2018).

58. S. Li, Y. Park, S. Duraisingham, F. H. Strobel, N. Khan, Q. A. Soltow, D. P. Jones, B. Pulendran, Predicting network activity from high throughput metabolomics. PLoS Comput Biol 9, e1003123 (2013).

59. R-Core-Team, A language and environment for statistical computing. R Foundation for Statistical Computing, Vienna, Austria.. (2019).

60. H. Wickham, ggplot2: Elegant Graphics for Data Analysis.. Springer-Verlag New York, (2016).

61. B. Yilmaz, M. R. Spalinger, L. Biedermann, Y. Franc, N. Fournier, J. B. Rossel, P. Juillerat, G. Rogler, A. J. Macpherson, M. Scharl, The presence of genetic risk variants within PTPN2 and PTPN22 is associated with intestinal microbiota alterations in Swiss IBD cohort patients. PloS one 13, e0199664 (2018).

62. A. S. Whiteley, S. Jenkins, I. Waite, N. Kresoje, H. Payne, B. Mullan, R. Allcock, A. O’Donnell, Microbial 16S rRNA Ion Tag and community metagenome sequencing using the Ion Torrent (PGM) Platform. J Microbiol Methods 91, 80-88 (2012).

63. B. Yilmaz, P. Juillerat, O. Øyås, C. Ramon, F. D. Bravo, Y. Franc, N. Fournier, P. Michetti, C. Mueller, M. Geuking, V. E. H. Pittet, M. H. Maillard, G. Rogler, R. Wiest, J. Stelling, A. J. Macpherson, S. I. C. Investigators, Microbial network disturbances in relapsing refractory Crohn’s disease. Nat Med 25, 323-336 (2019).

64. P. J. McMurdie, S. Holmes, Phyloseq: a bioconductor package for handling and analysis of high-throughput phylogenetic sequence data. Pac Symp Biocomput, 235-246 (2012).

65. B. J. Callahan, K. Sankaran, J. A. Fukuyama, P. J. McMurdie, S. P. Holmes, Bioconductor Workflow for Microbiome Data Analysis: from raw reads to community analyses. F1000Res 5, 1492 (2016).

66. X. C. Morgan, T. L. Tickle, H. Sokol, D. Gevers, K. L. Devaney, D. V. Ward, J. A. Reyes, S. A. Shah, N. LeLeiko, S. B. Snapper, A. Bousvaros, J. Korzenik, B. E. Sands, R. J. Xavier, C. Huttenhower, Dysfunction of the intestinal microbiome in inflammatory bowel disease and treatment. Genome Biol 13, R79 (2012).

67. F. Asnicar, G. Weingart, T. L. Tickle, C. Huttenhower, N. Segata, Compact graphical representation of phylogenetic data and metadata with GraPhlAn. PeerJ 3, e1029 (2015).

68. M. Ionac, One technique, two approaches, and results: thoracic duct cannulation in small laboratory animals. Microsurgery 23, 239-245 (2003).

69. H. Y. Benjamini Y., Controlling the false discovery rate: a practical and powerful approach to multiple testing. J. R. Statist. Soc 57, 289-300 (1995).

70. D. Szklarczyk, A. Franceschini, S. Wyder, K. Forslund, D. Heller, J. Huerta-Cepas, M. Simonovic, A. Roth, A. Santos, K. P. Tsafou, M. Kuhn, P. Bork, L. J. Jensen, C. von Mering, STRING v10: protein-protein interaction networks, integrated over the tree of life. Nucleic acids research 43, D447-452 (2015).

